# Functional characterization of luciferase in a brittle star indicates parallel evolution influenced by genomic availability of haloalkane dehalogenase

**DOI:** 10.1101/2024.10.14.618359

**Authors:** Emily S Lau, Marika Majerova, Nicholai M Hensley, Arnab Mukherjee, Michal Vasina, Daniel Pluskal, Jiri Damborsky, Zbynek Prokop, Jérôme Delroisse, Wendy-Shirley Bayaert, Elise Parey, Paola Oliveri, Ferdinand Marletaz, Martin Marek, Todd H Oakley

## Abstract

Determining why convergent traits use distinct versus shared genetic components is crucial for understanding how evolutionary processes generate and sustain biodiversity. However, the factors dictating the genetic underpinnings of convergent traits remain incompletely understood. Here, we use heterologous protein expression, biochemical assays, and phylogenetic analyses to confirm the origin of a luciferase gene from haloalkane dehalogenases in the brittle star *Amphiura filiformis*. Through database searches and gene tree analyses, we also show a complex pattern of presence and absence of haloalkane dehalogenases across organismal genomes. These results first confirm parallel evolution across a vast phylogenetic distance, because octocorals like *Renilla* also use luciferase derived from haloalkane dehalogenases. This parallel evolution is surprising, even though previously hypothesized, because many organisms that also use coelenterazine as the bioluminescence substrate evolved completely distinct luciferases. The inability to detect haloalkane dehalogenases in the genomes of several bioluminescent groups suggests that the distribution of this gene family influences its recruitment as a luciferase. Together, our findings highlight how biochemical function and genomic availability help determine whether distinct or shared genetic components are used during the convergent evolution of traits like bioluminescence.

## Introduction

Similar traits evolve convergently using shared or distinct genetic pathways, depending on the interplay between function, mutation, and phylogenetic history (Christin et al. 2010; Stern 2013). Similar traits may originate repeatedly via parallel evolution in distinct lineages by recruiting homologous genes; especially when genetic pathways are shared among lineages (Shubin et al. 2009; Rosenblum et al. 2014), or when functional evolution is constrained by limited genetic solutions (Lau et al. 2024). Conversely, similar traits may originate repeatedly by using distinct and non-homologous genes if there are many possible genetic pathways that produce the same function (Tomarev and Piatigorsky 1996; Foster et al. 2022) or if shared genetic pathways are not maintained (Oakley 2024). Here, we explore the factors shaping the repeated evolution of coelenterazine-based bioluminescence.

Bioluminescence, the production of light by a living organism, is an excellent system for studying patterns of convergence. Bioluminescence repeatedly evolved at least 94 times across distantly related taxa (Lau and Oakley 2020) and is produced when enzymes, generally called luciferases, oxidize any of a number of substrates generally called luciferins (Shimomura 2019). Across convergent origins of bioluminescence, many luciferases are non-homologous and taxon specific, whereas the same luciferin may be used in many bioluminescence systems, even across vast phylogenetic distances (Delroisse et al. 2021). The most widespread luciferin in marine bioluminescence systems is called coelenterazine, which is produced by a few taxa, such as the shrimp *Systellaspis debilis* (Thomson et al. 1995), the copepod *Metridia pacifica* (Oba et al. 2009), and the ctenophores *Mnemiopsis leidyi* and *Bolinopsis infundibulum* (Bessho-Uehara et al. 2020). Other luminous organisms, such as the jellyfish *Aequorea* (Haddock et al. 2001), the shrimp *Gnathophausia ingens* (Frank et al. 1984), and the brittle star *Amphiura filiformis* (Mallefet et al. 2020), obtain coelenterazine through their diets. Despite using the same luciferin, most organisms that use coelenterazine evolved luciferases by recruiting non-homologous genes (Markova and Vysotski 2015), revealing a diversity of genetic solutions for coelenterazine-based light production. This previous work suggests that coelenterazine-based bioluminescence typically evolves convergently, rather than in parallel.

Surprisingly, sea pansies and brittle stars may have repeatedly recruited members of the haloalkane dehalogenase gene family to be coelenterazine-based luciferases (Delroisse et al. 2017; Chaloupkova et al. 2019). The sea pansies *Renilla* sp. use a luciferase that was first cloned in 1991 (Lorenz et al. 1991) and has been structurally (Loening et al. 2007) and biochemically (Schenkmayerova et al. 2023) well-characterized. The brittle star *Amphiura filiformis* may use a luciferase homologous to haloalkane dehalogenases, based on the immunohistochemical detection of *Renilla* luciferase-like proteins in the light-emitting spines of their arms (Delroisse et al. 2017). However, while several candidate genes were identified from the genome of *A. filiformis* (Delroisse et al. 2017; Parey et al. 2024), the luciferase gene has not yet been identified and biochemically characterized. Determining whether these distantly related taxa share a common biochemical mechanism — and if so, understanding the processes that shape the repeated recruitment of this gene family during the evolution of coelenterazine-based bioluminescence — requires identifying the luciferase gene of *A. filiformis* and investigating the distribution of this gene family across luminous organisms.

We recombinantly expressed and functionally tested haloalkane dehalogenase/luciferase (hereafter HLD/LUC) genes from the genome of *A. filiformis* and identified one HLD/LUC gene, which we name *Amphiura* luciferase, or “*afLuc*”, encoding a protein with robust luciferase activity. We also identified a gene, which we named *Amphiura filiformis* dehalogenase, or “*dafA*”, encoding a protein with dehalogenase activity and low luciferase activity. Similar to *Renilla* luciferase (RLuc), AfLuc lacks dehalogenase activity with a common substrate, 1,2-dibromoethane, while DafA exhibits activity with 1,2-dibromoethane and other halogenated compounds. AfLuc produces luminescence with an emission spectrum similar to RLuc’s, with maximum light emission at a wavelength of 482 nm, and exhibits a similar affinity for coelenterazine. Haloalkane dehalogenase genes in metazoans may have originated via a horizontal gene transfer from bacteria to a cnidarian-bilaterian ancestor (Delroisse et al. 2017) and subsequent gene losses may have influenced the availability of this gene family for recruitment during the evolution of bioluminescence. Altogether, our results provide functional evidence for the evolution of luciferases in brittle stars in parallel with sea pansies, a finding that deviates from the typical pattern of convergent genetic recruitment in coelenterazine-based systems, highlighting how genetic processes such as horizontal gene transfer and gene loss impact the predictability of convergent evolution and subsequent biodiversity.

## Materials and methods

For a comprehensive list of materials and more detailed methods used in this study, please refer to the methods in the supplemental material.

### Obtaining gene sequence and expression data

We obtained HLD/LUC sequences from *Amphiura filiformis* from various sources, as follows. We obtained the sequences Gen224433 and Gen313061 (named *afLuc*) from a previous transcriptomic dataset (Delroisse et al. 2017), and the sequence Uni20302.6 from an initial *de novo* transcriptome of the species (Delroisse et al. 2014). The sequences AF10707.1, AF17859.1, AF37282.1, AF37308.1 (named *dafA*), and AF37332.1 originated from a set of preliminary gene models predicted from a genome of *A. filiformis* — new versions of these gene models are now published in Parey et al. (2024). Based on percent sequence identity, we synonymized all sequences from the transcriptomic dataset, preliminary gene models, and final gene models from Parey et al. (2024) (Supplemental Table S1). Additionally, we obtained the gene expression dataset used in this study from Parey et al. (2024), which combined expression data from several publications (Delroisse et al. 2014; Delroisse et al. 2015; Dylus et al. 2016).

### Primer design, A. filiformis sampling, genomic DNA extraction and amplification of dehalogenase sequences

We performed genomic DNA-based validation PCRs to confirm portions of the HLD/LUC gene sequences (Supplemental Figure S1). For each gene, we designed primer pairs using the Primer3 software (v4.1.0, http://bioinfo.ut.ee/primer3) (Supplemental Table S2). We collected *A. filiformis* individuals from a depth of 30-40 meters in the Gullmars fjord near the Kristineberg Marine Research Station (University of Gothenburg, Fiskebäckskil, Sweden) and extracted genomic DNA from arm tissues using Qiagen DNeasy® Blood & Tissue kit. We performed PCR amplifications using Red’y’Star Mix (Eurogentecs) or Q5® High-Fidelity DNA Polymerase (New England BioLabs) and purified PCR products prior to sending samples for Sanger sequencing (Eurofins Genomics, Germany). We aligned these sequences with the reference HLD/LUC genes to verify their identities (Supplemental Figure S2). For more detailed protocols for DNA extractions and PCR, please refer to the methods in the supplemental materials.

### Expressing recombinant proteins and testing crude cellular extracts for luciferase activity

We codon-optimized and synthesized DNA sequences corresponding to the luciferase sequence of *Renilla reniformis* (UniProt Accession P27652) and dehalogenase sequences from *Amphiura filiformis*, namely Gen224433, Gen313061 (named *afLuc*), and Uni20302.6 as reported in (Delroisse et al. 2017), and AF10707.1, AF17859.1, AF37282.1, AF37308.1 (named *dafA*), AF37332.1 (sequences predicted from a draft genome of *A. filiformis*), and the pyrosome luciferase (*pyroLuc*), identified by Tessler et al. (2020). We cloned these sequences into the bacterial expression vector pET21b, transformed competent *E. coli* cells for propagating plasmids, then extracted and used Sanger sequencing to confirm successful cloning. Then, we transformed competent BL21 cells via electroporation with these plasmids for protein expression. We grew up transformed BL21 cells in Terrific Broth containing ampicillin, at 37 °C and shaking at 250 rpm, until cultures reached mid-log phase. We then added isopropyl-β-D-thiogalactopyranoside (IPTG) to induce protein expression, moved the cultures to a shaker at room temperature, and continued protein expression for 16-18 hours. We centrifuged the cultures to harvest bacterial cells, removed the supernatant, and froze the cell pellets. We lysed bacterial cells in a lysis buffer and sonicated the cells on ice. Then, we centrifuged the lysed cells and collected the supernatant, which contained our recombinant proteins. We tested the clarified supernatant from lysed bacterial cells for luciferase activity by adding coelenterazine and measuring luminescence using a microplate reader.

### Recombinant expression and purification of AfLuc, RLuc, DafA, and PyroLuc using immobilized metal chelate affinity chromatography

We expressed recombinant proteins using the protocol as described above. After harvesting and freezing cell pellets, we extracted recombinant proteins by resuspending bacterial cells in lysis buffer containing imidazole and sonicating the cells on ice. We centrifuged these lysates to pellet cellular debris, collected the clarified supernatants, added it to Ni-NTA agarose beads, then incubated the samples at 4 °C overnight while mixing on a rotary mixer. Next, we loaded the Ni-NTA slurry into gravity-flow chromatography columns, discarded the flow through, washed the column twice with wash buffer, and eluted proteins bound to agarose using an elution buffer. We performed spin ultrafiltration (10 kDa molecular weight cutoff) to concentrate and buffer exchange the eluates into a storage buffer. After running SDS-PAGE to assess protein purity and quantifying proteins via a Bradford assay, we flash froze single-use aliquots of recombinant protein and stored them in −80 °C until use.

### Estimating Michaelis-Menten kinetic profiles and luminescence decay parameters

We characterized the enzyme kinetics of AfLuc, DafA, and PyroLuc, using *Renilla* luciferase (RLuc) as a positive control. Since coelenterazine may produce low amounts of chemiluminescence with proteins such as bovine serum albumin (BSA) (Vassel et al. 2012), we used BSA as a negative control. We added varying concentrations of coelenterazine to recombinant protein (final concentrations 10 μM, 5 μM, 2.5 μM, 1.25 μM, 0.625 μM, 0.3125 μM, 0.15625 μM) and measured light production. Specifically, we used a plate reader (Tecan Spark) to measure the background luminescence for five cycles prior to injecting coelenterazine and measuring luminescence for 30 cycles, with a 1 second integration time for each cycle. We repeated each sample measurement in triplicate, and for each measurement, we subtracted the background luminescence. For detailed methods for kinetic data analysis and statistics, please refer to the Supplemental Methods section.

We used nonlinear model fitting and comparison to estimate parameters describing the decay of light production between different proteins measured in the plate reader, as above. Identified from the literature, we fit 4 different models of exponential decay separately to each dataset in R using nlsLM, which uses the more robust LM method to find suitable parameter estimates. For each protein, we then compared models using the corrected Akaike Information Criteria; the model with the lowest value was considered the best fit. However, we note that for one sample (AfLuc) some models were less than 2 AICc values apart, indicating model equivalency. We also visually examined the predicted fit of every model to the data in each dataset.

### Luminescence emission spectra measurements

We measured emission spectra using a custom spectroradiometer set-up at UCSB, as detailed in a previous study (Hensley et al. 2021). In brief, we added coelenterazine to recombinant proteins diluted with 1X Tris Buffered Saline (TBS), and measured the emission spectra using a spectroradiometer (Acton SpectraPro 300i) with a charge-coupled device camera detector (Andor iDus). We corrected these spectral data using correction factors calculated from the spectrum of a black body-like light source (Ocean Optics LS-1) and subtracted background emission spectra data of 1X TBS from the experimental data. We repeated each sample measurement in triplicate, then normalized and averaged these data.

### Testing whole cell extracts for dehalogenase activity

We prepared whole cell extracts by transferring transformed BL21 cells into sterile 96-well plates, then incubated the plates for 3 hours at 37 °C while shaking at 200 rpm. We added IPTG to induce protein expression to each well and incubated the plates at 20 °C for 18 hours while at 200 rpm. We centrifuged the 96-well plates to pellet cell cultures, washed the pellets with reaction buffer twice, and centrifuged again to harvest cell pellets. We resuspended cell pellets in the reaction buffer and lysed them by freezing them at −70 °C.

To screen whole cell extracts for dehalogenase activity, we used a halide oxidation (HOX) assay (Aslan-Üzel et al. 2020). In a new 96-well plate, we added the assay master mix, which consists of 25 µM aminophenyl fluorescein, 26 mM H_2_O_2_, 1.1 U *Curvularia inaequalis* histidine-tagged vanadium chloroperoxidase, 1 mM orthovanadate, 20 mM phosphate buffer, pH = 8.0. We then added resuspended cells to each well for a final OD600 ∼ 0.02 and added the 1,2-dibromoethane (DBE) substrate. We measured fluorescence with an excitation at 488 nm and emission detention at 525 nm, at 30 °C (Synergy™ H4 Hybrid Microplate Reader), with results normalized to OD600 = 1. We measured all data in triplicate and calculated means and standard deviations.

### Measurements of specific dehalogenase activity at varying temperatures

We measured the temperature profile and dehalogenase activity with various substrates for DafA using the capillary-based droplet microfluidic platform MicroPEX (Buryska et al. 2019; Vasina et al. 2022), which enables us to measure specific enzyme activity within droplets for multiple enzyme variants in one run. Briefly, we generated a custom sequence of droplets (Mitos Dropix), then incubated these droplets with the halogenated substrate in a reaction solution and a complementary fluorescent indicator, 8-hydroxypyrene-1,3,6-trisulfonic acid. Then, fluorescence was measured using an optical setup with an excitation laser (450 nm), a dichroic mirror with a cut-off at 490 nm filtering the excitation light, and a Si-detector. We processed raw data using LabView and used MatLab to calculate specific activities.

### Phylogenetic analysis of dehalogenase sequences

Using AfLuc (Accession PP777633), RLuc (Accession P27652), PyroLuc (Accession PP777641), DhaA (Accession P59336), LinB (Accession D4Z2G1), DhlA (Accession P22643), and DrbA (Accession G3XCP3) as query sequences, we used DIAMOND blastp (v 0.9.12.113) (Buchfink et al. 2015) to identify the top 50 proteins with the lowest e-value scores from the UniRef90 database (downloaded March 2024). We used HMMER (v 3.4) to identify alpha/beta hydrolase domains (PF00561) present in all haloalkane dehalogenases and aligned these domain sequences using MAFFT (v 7.453) (Katoh and Standley 2013). We used IQ-TREE (v 2.0.3) (Minh et al. 2020) to infer a maximum likelihood phylogeny using the best-fit substitution model (LG + R7) as determined by ModelFinder (Kalyaanamoorthy et al. 2017) according to Bayesian Information Criterion and performed ultrafast bootstrap approximation with 1000 replicates. We visualized trees using iToL and annotated the phylogeny based on the taxon ID of the representative sequence for each UniRef90 accession.

## Results

### The genome of A. filiformis encodes a gene with high luciferase activity

We identified one gene, *afLuc*, which encodes a protein with luciferase activity but no dehalogenase activity, and one gene, *dafA*, which encodes a protein with dehalogenase activity and low luciferase activity in *A. filiformis*. The gene *afLuc* corresponds to two previously identified gene models (Gen313061 and AFI20122.1) and the gene *dafA* corresponds to two previously identified gene models (AF37308.1 and AFI06958.1). Based on a luciferase assay, which measures light production upon addition of coelenterazine (Figure 1A, top), AfLuc exhibited statistically significant luciferase activity (Dunnett’s test, p-value < 2e^−16^), compared to a negative control of bovine serum albumin (BSA). DafA exhibits low luciferase activity that was significantly higher than the negative control (Dunnett’s test, p-value < 2e^−16^), but produced light four and five orders of magnitude lower than that of AfLuc and *Renilla* luciferase (RLuc), respectively (Figure 1B, Supplemental Figure S3).

**Figure 1.**
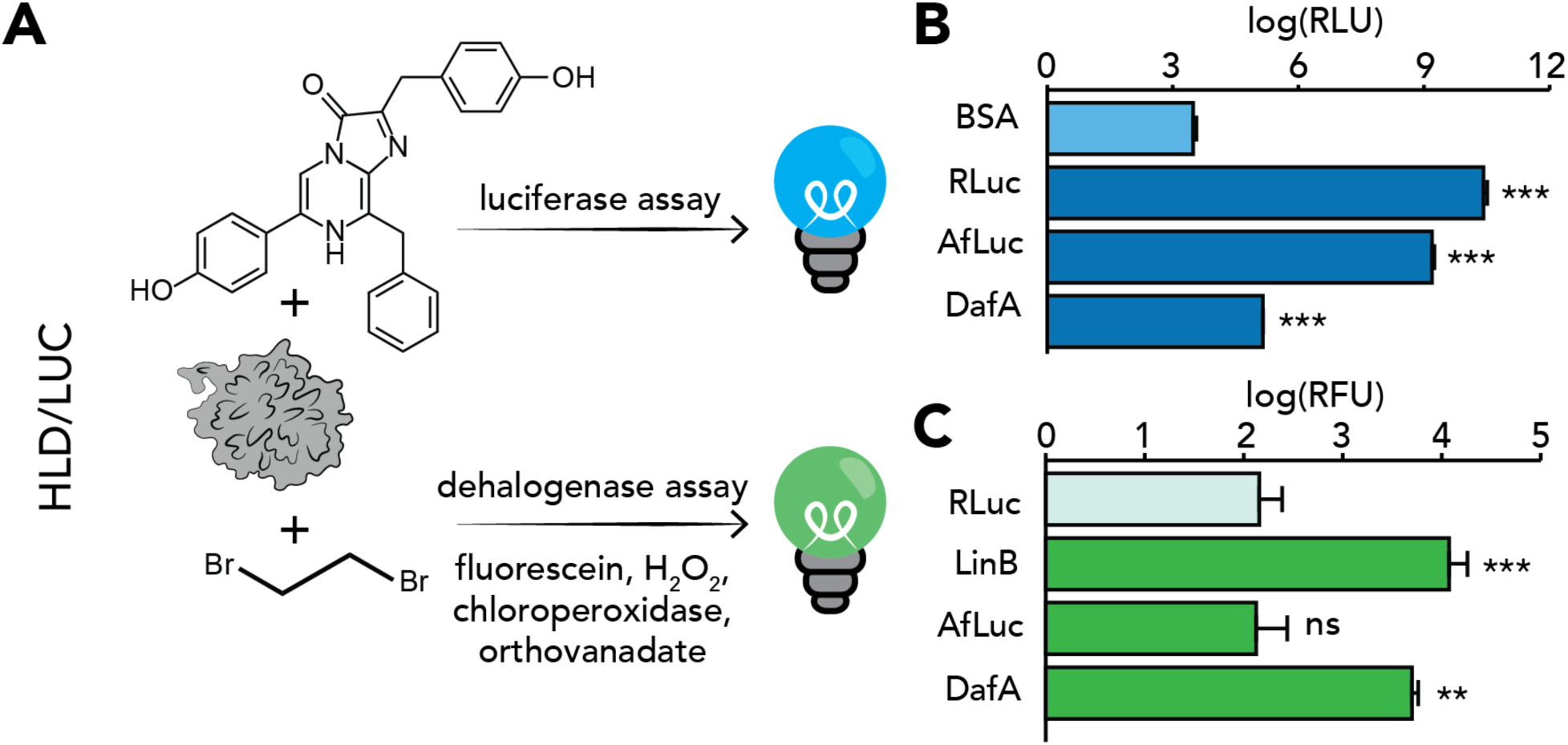
*Amphiura* luciferase (AfLuc) and DafA exhibit significant luciferase activity, but only DafA exhibits significant dehalogenase activity. (A) We tested HLD/LUC proteins for luciferase activity with coelenterazine substrate and dehalogenase activity with 1-2-dibromoethane substrate. (B) Results of luciferase activity reveal AfLuc and DafA exhibit significant luciferase activity when compared to the negative control of BSA. Luciferase activity is quantified in relative light units (RLU). The positive control is *Renilla* luciferase (RLuc) and the negative control is bovine serum albumin (BSA, light blue). (C) Dehalogenase assay reveals DafA, but not AfLuc or other HLD/LUC proteins tested (Supplemental Figures S4 and S5), exhibits dehalogenase activity with 1-2-dibromoethane as a substrate. Dehalogenase activity is quantified in relative fluorescence units (RFU). We compared dehalogenase activity at 60 minutes for AfLuc and DafA, using LinB as a positive control and RLuc as a negative control (light green). Data in this figure are expressed as average ± standard deviation represented by the error bars (N = 3). * denotes P < 0.05, ** denotes P < 0.01, *** denotes P < 0.001, and ns denotes non-significance (P ≥ 0.05)

Based on a halide oxidation (HOX) assay (Figure 1A, bottom), a fluorescence-based assay that quantifies dehalogenase activity, only DafA exhibited statistically significant dehalogenase activity (Dunnett’s test, p-value = 0.00402) with the substrate 1,2-dibromoethane (Figure 1C, Supplemental Figure S4 and S5). Further functional tests revealed that DafA exhibits maximum activity toward 1,2-dibromoethane at 30 °C (Supplemental Figure S6), and at this temperature, it also catalyzed the dehalogenation of 1-iodohexane, 1,3-dibromopropane, 1-bromo-3-chloropropane, and 3-chloro-2-methylpropene (Supplemental Figure S7). The highest specific activity was measured towards 1,2-dibromoethane (74.8 ± 0.8 nmol s^−1^ mg^−1^), while the lowest activity was measured towards 1-iodohexane (6.5 ± 1.5 nmol s^−1^ mg^−1^). The activity of DafA is comparable to those of characterized haloalkane dehalogenases in bacteria (Supplemental Table S3).

### RLuc and AfLuc exhibit similar emission spectra and catalytic properties

We conducted a conventional biochemical characterization (Figure 2A and 2C, Supplemental Figure S8) followed by a global numerical analysis (Johnson 2019) incorporating new standards for the collection and fitting of steady-state kinetic data (Supplemental Figure S9 and Supplemental Table S4). Unlike the traditional analysis of initial velocity, the updated numerical approach enables direct estimation of the turnover number *k*_cat_ without requiring complex luminometer calibration or quantum yield determination (Schenkmayerova et al. 2021).

**Figure 2.**
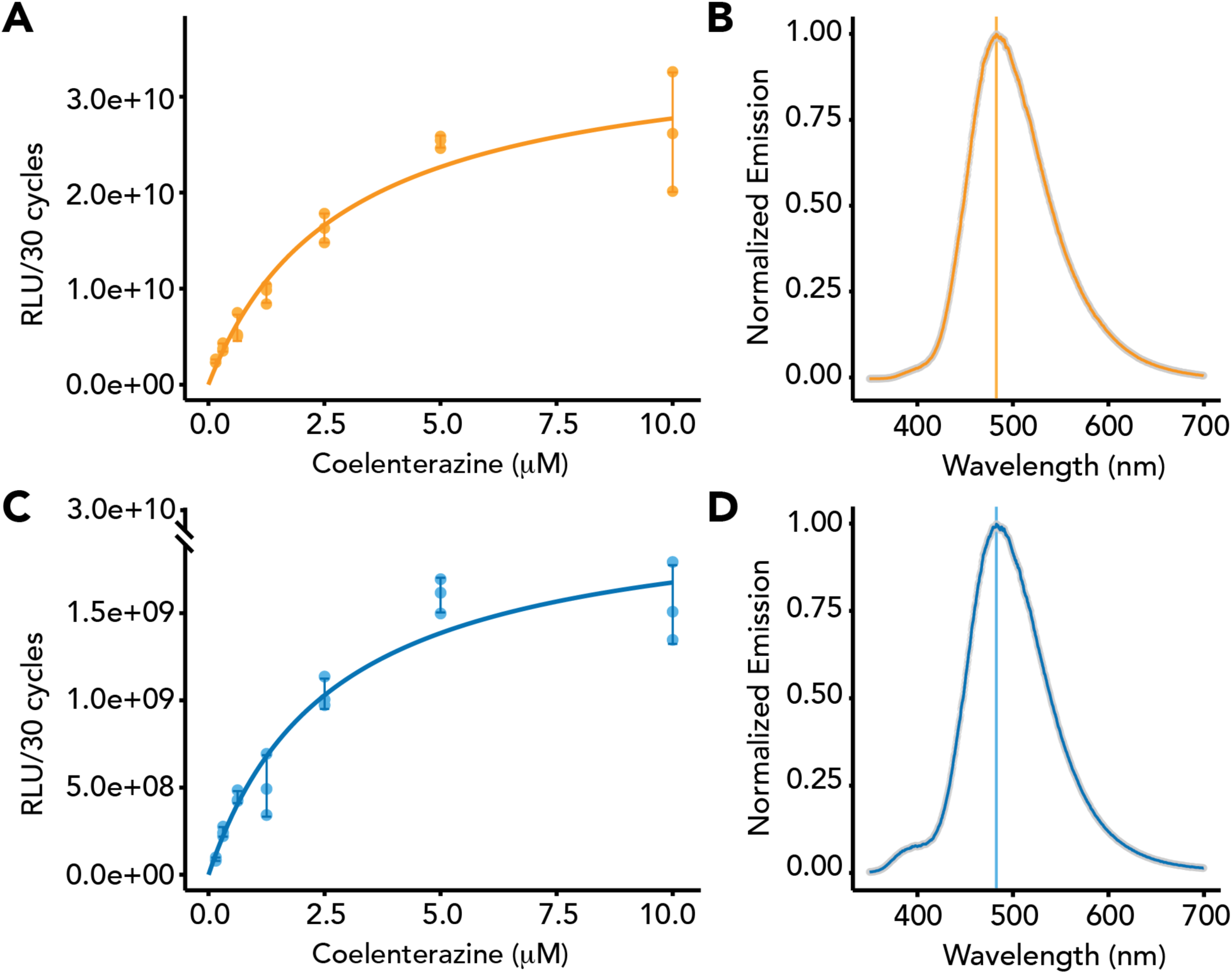
Biochemical properties of RLuc (orange) and AfLuc (blue). (A, C) Steady-state kinetic data recorded upon mixing 25 nM of protein with varying concentrations of coelenterazine. We averaged these data (N = 3) and fit them to a Michaelis-Menten model. Data are expressed as average ± standard deviation represented by the error bars (N = 3). (B, D) RLuc and AfLuc emit bioluminescence with similar maximum wavelengths and have similar emission spectra, but AfLuc’s emission spectrum has a small shoulder at around 400 nm.

The kinetic analysis indicates that AfLuc has kinetic parameters comparable to those of RLuc, with substrate affinity *K*_m_ = 1.21 ± 0.03 µM and *k*_cat_ = 4.12 ± 0.05 s^−1^ for AfLuc, and *K*_m_ = 0.91 ± 0.05 µM and *k*_cat_ = 4.2 ± 0.2 s^−1^ for RLuc. The kinetic parameters of RLuc are consistent with previously reported values of *K*_m_ = 1.5 ± 0.1 µM and *k*_cat_ 4.7 ± 0.1 s^−1^, as determined using the updated protocol for collecting and fitting steady-state kinetic data (see Supplemental Methods). Interestingly, the global kinetic analysis further indicated that AfLuc does not undergo the irreversible inactivation observed in RLuc and other tested variants. The absence of inactivation is also clearly visible from the luminescence decay data (Supplemental Figure S10). In this analysis, AfLuc is the only variant displaying a consistent single-exponential decay of luminescence activity over time, whereas the other variants demonstrate a significant slowdown in kinetics, characterized by a biexponential decay model (Supplemental Table S5).

Both AfLuc and RLuc exhibit similar emission spectra (RLuc λ_max_ = 482.20 nm and FWHM = 91.92, AfLuc λ_max_ = 482.93 nm and FWHM = 91.77). However, one difference is that AfLuc’s emission spectrum shows a minor shoulder around 400 nm (Figure 2B and D).

### AfLuc is mainly expressed in adult tissues while DafA is expressed throughout development

The gene *afLuc* appears to be the most highly expressed HLD/LUC gene in the adult arm tissue, but has little to no expression during other developmental timepoints (Figure 3). Only *dafA*, and to a much lesser extent the gene models AFI17177.1 and AFI14276.1, are expressed during development. We observed that closely related HLD/LUC genes exhibit similar gene expression patterns across development. For instance, AFI21141.1, AFI19872.1, AFI20122.1/*afLuc*, and AFI19881.1 are highly expressed primarily in the adult arms, while AFI06721.1 and AFI06853.1 are lowly expressed only during the early developmental stages.

**Figure 3.**
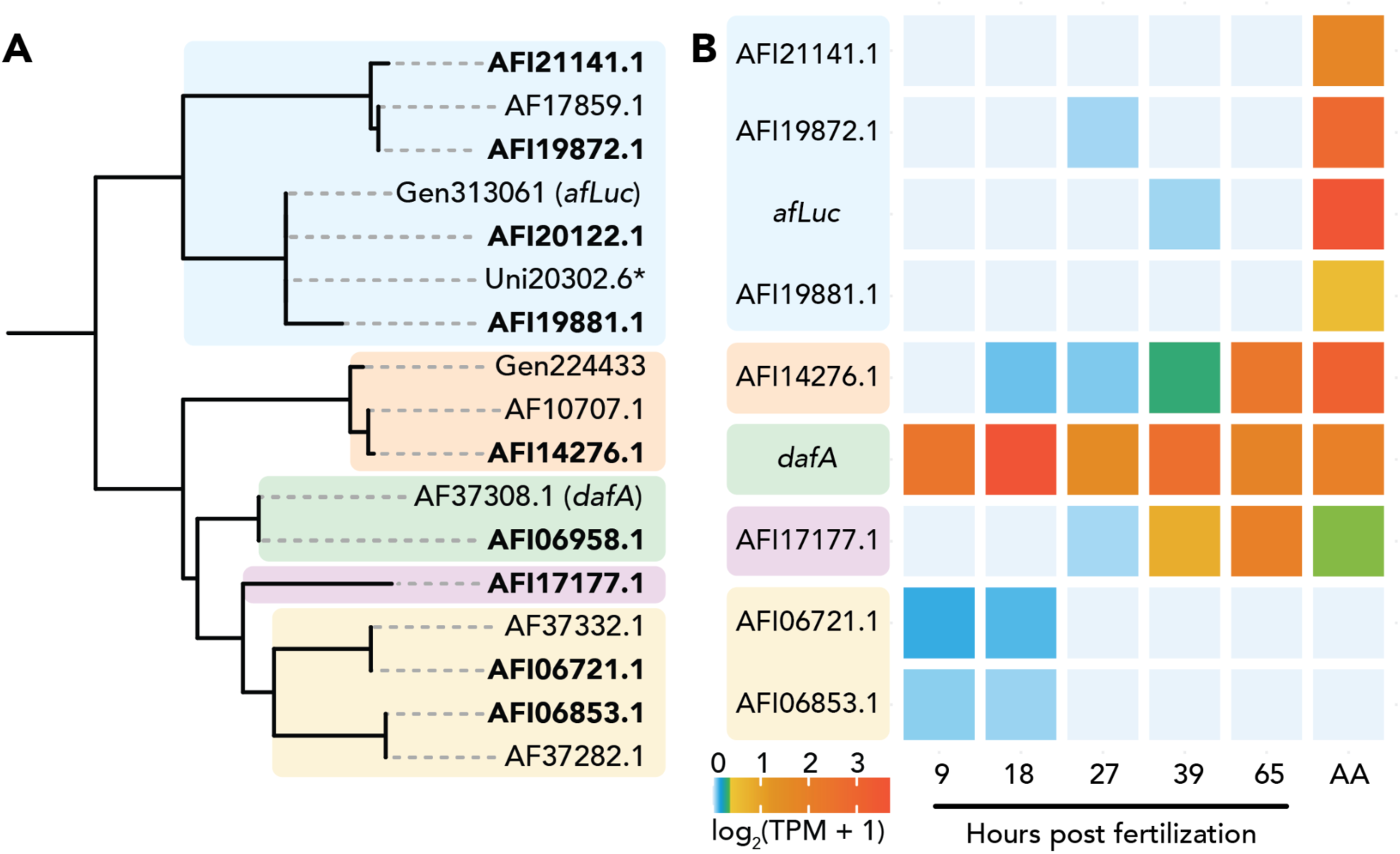
Closely related HLD/LUC genes exhibit similar expression patterns during development. (A) Maximum likelihood, midpoint rooted phylogeny of HLD/LUC protein sequences from preliminary gene models and the final gene models (bolded) as published in Parey et al. (2024). * Uni20302.6 is a transcript sequence derived from a transcriptome of the adult arm, and encodes a truncated protein sequence, which is otherwise identical to the protein sequence encoded by Gen313061. (B) Heatmap showing gene expression of HLD/LUC genes during development and in the adult arm (AA). Gene expression dataset is from a publication (Parey et al. 2024), which quantified gene expression levels in log_2_(Transcripts Per Million (TPM) + 1). During the development of *Amphiura filiformis*, luminescence ability emerges after larvae metamorphose into juveniles. The gene *afLuc* is most highly expressed in the arms of adult specimens, where bioluminescence is produced. The gene *dafA* is highly expressed during earlier developmental time points and in the adult arms.

### Amphiura filiformis and Renilla catalyze light production by using homologous genes

Dehalogenase genes in *Renilla* and *A. filiformis* independently evolved luciferase activity with coelenterazine. They may have evolved from a haloalkane dehalogenase gene family that originally was horizontally transferred from bacteria to metazoans (Figure 4). We identified dehalogenase-like proteins — containing conserved alpha-beta hydrolase domains (Chovancová et al. 2007) — in bacteria and eukaryotes, namely Fungi, Porifera, Cnidaria, Annelida, Hemichordata, Echinodermata, and Chordata. Our phylogenetic analysis identified three distinct clades of alpha-beta hydrolases originating from dehalogenases, each clade containing at least one representative bacterial dehalogenase from subfamilies HLD-I, HLD-II, and HLD-III (Chovancová et al. 2007).

**Figure 4.**
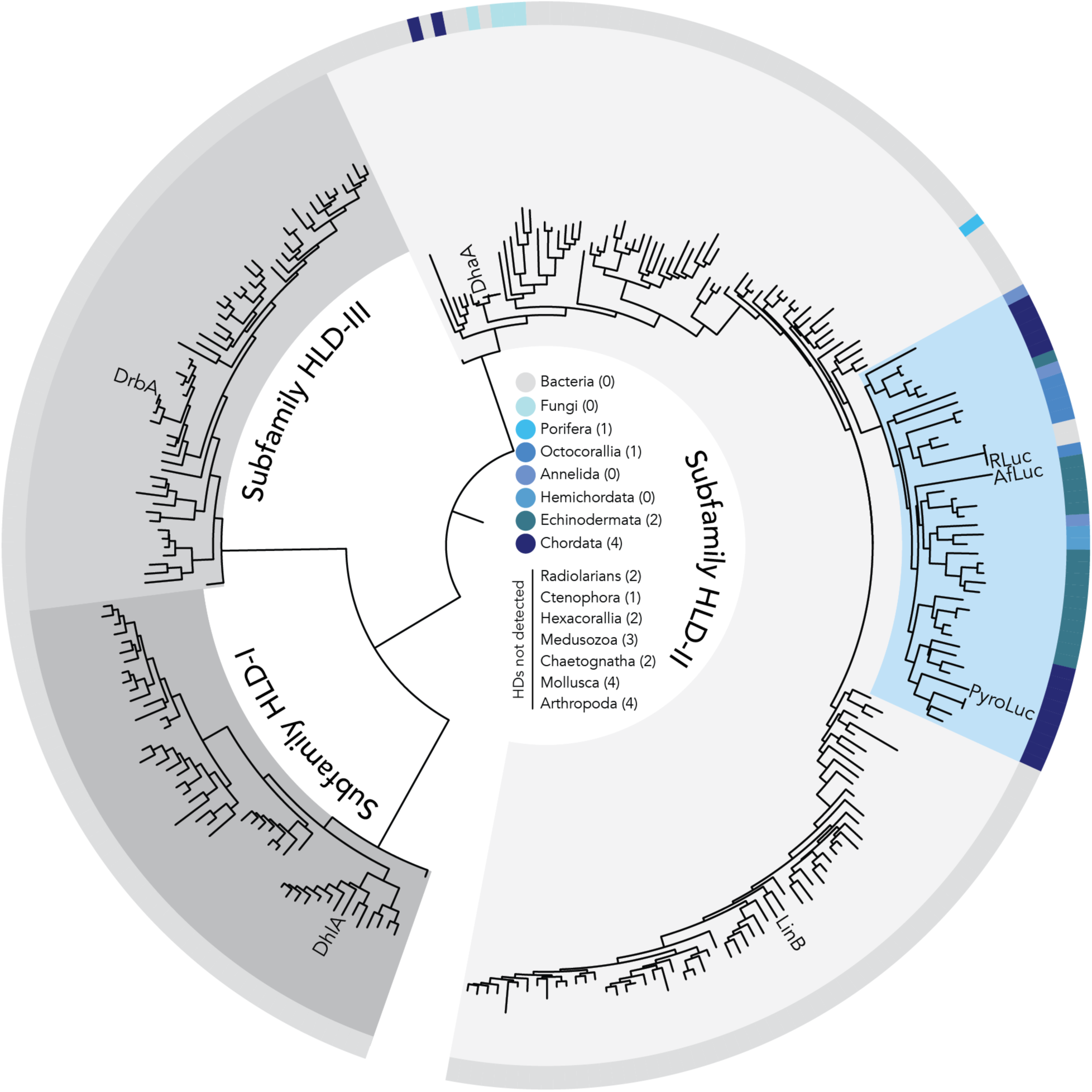
Maximum likelihood phylogeny of alpha-beta hydrolase domains from haloalkane dehalogenase-like sequences. We used characterized haloalkane dehalogenase sequences from bacteria (DhlA, DrbA, DhaA, and LinB) and luciferases (RLuc, AfLuc, and PyroLuc) as query sequences to identify haloalkane dehalogenase-like proteins in the UniRef90 database. Clades of dehalogenase subfamilies are colored in shades of gray. All metazoan sequences are found within subfamily HLD-II and most sequences are found in one clade (blue), supporting a horizontal gene transfer from bacteria to a cnidarian-bilaterian ancestor. The outer arc is colored based on taxonomy (gray = bacteria, shades of blue = eukaryotes), as denoted in the legend found in the center of the tree. Numbers next to each taxon name indicate the number of times coelenterazine-based bioluminescence repeatedly evolved (Supplemental Table S6).

All eukaryotic sequences are found within a clade containing sequences from bacterial dehalogenases in subfamily HLD-II. Within this clade, fungal and metazoan sequences are polyphyletic, which suggests multiple horizontal transfers of a subfamily II dehalogenase gene from bacteria to eukaryotes. AfLuc is found within a clade of sequences from non-luminous metazoans, while RLuc is found in a clade of sequences from bacteria and non-luminous octocorals. Altogether, these results support the parallel evolution of luciferases in *Renilla* and *Amphiura filiformis* from a subfamily HLD-II dehalogenases, which may have originated in metazoans from an ancient bacterial horizontal gene transfer (Figure 4).

## Discussion

Convergently evolved traits may recruit homologous or non-homologous genes, depending on the range of possible genetic solutions and the availability of raw genetic material. In this study, we provide functional evidence supporting the parallel evolution of luciferases in sea pansies and brittle stars. While the genome of *A. filiformis* encodes multiple genes homologous to the luciferase from the sea pansy *Renilla* and other members of the haloalkane dehalogenase gene family, we identify only one gene encoding a functional luciferase. We present several lines of evidence supporting that *afLuc* is a functional luciferase gene in the bioluminescence system of *A. filiformis*. First, the light produced by AfLuc is strongly detectable and four orders of magnitude higher than that of DafA, a dual-function enzyme we find to have dehalogenase activity and low luciferase activity. Interestingly, the same pattern of dual activity was achieved by site-directed mutagenesis of a single active-site residue of Rluc (Chaloupkova et al. 2019) and by ancestral sequence reconstruction of dehalogenase and luciferase sequences (Schenkmayerova et al. 2021). Second, *afLuc* is highly expressed in *A. filiformis*’s arms, which produce bioluminescence and are where *Renilla* luciferase-like proteins are localized (Delroisse et al. 2017). While *afLuc* has little to no expression during early developmental stages, it shows low expression during early stages of arm regeneration and strong expression during the late stages of regeneration (Parey et al. 2024). Consistent with this expression pattern, luciferase activity is detected in juvenile *A. filiformis* only after their arms start to develop (Coubris et al. 2024). Third, similar to RLuc, AfLuc does not exhibit dehalogenase activity with the substrate 1,2-dibromoethane, supporting a shift from dehalogenase to luciferase function. Taken together, these lines of evidence strongly support AfLuc’s organismal role in the bioluminescence system of *A. filiformis*.

Phylogenetic analysis of dehalogenase sequences supports the parallel evolution of luciferases in *A. filiformis* and *Renilla*. The phylogenetic distribution of dehalogenase genes is widespread in bacteria (Janssen et al. 2005) but sparse in fungi and metazoans. Our phylogeny, which contains representative sequences from the three bacterial haloalkane dehalogenase subfamilies (Chovancová et al. 2007) and similar sequences in eukaryotes, suggests that the conserved alpha-beta hydrolase domains from haloalkane dehalogenase genes were horizontally transferred, multiple times, from bacteria to eukaryotes. Additionally, the sequence of the alpha-beta hydrolase domain in RLuc is identical to the hydrolase domain found in the bacterial cluster UniRef90_A0A941CXK5, with a representative aminoglycoside phosphotransferase sequence from the bacteria *Allobacillus saliphilus*, which supports a secondary transfer of the hydrolase domain from *Renilla* to bacteria. In bacteria, dehalogenases are often associated with molecules implicated in the transfer of genetic material (e.g., integrase and invertase genes, insertion elements), which implicates horizontal transfer as a mechanism for genetic recruitment in the evolution of xenobiotic degradation (Janssen et al. 2005). The presence of conserved alpha-beta hydrolase domains from bacterial dehalogenases in multi-domain proteins in prokaryotes and eukaryotes suggests that horizontal transfer may be an important genetic mechanism in the evolution of various functions, even besides light production and xenobiotic degradation. Overall, our phylogenetic results support an origin of haloalkane dehalogenase genes in metazoan genomes via a horizontal gene transfer from bacteria to an early cnidarian-bilaterian ancestor — as previously hypothesized by Delroisse et al. (2017) — and reveal that this gene family may have been subsequently lost in many metazoan lineages, including those with taxa that produce coelenterazine-based bioluminescence (Supplemental Table S6). Other possible explanations for the limited distribution of this gene family in Metazoa involve an initial horizontal gene transfer from bacteria to a metazoan, followed by multiple metazoan-to-metazoan horizontal gene transfers. Nevertheless, the limited availability of this gene family, coupled with the numerous alternative genetic solutions that can converge to produce coelenterazine-based bioluminescence (Lau and Oakley 2020), may help explain why haloalkane dehalogenases have not been more frequently recruited in the multiple evolutionary origins of this trait.

In addition to octocoral cnidarians and the brittle star *A. filiformis*, the bioluminescence system of the chordate *Pyrosoma atlanticum* may also use a luciferase homologous to haloalkane dehalogenases (Tessler et al. 2020). However, its usage in the pyrosome bioluminescence system remains suspect for several reasons. Primarily, the study that identified the pyrosome luciferase did not demonstrate the presence of coelenterazine *in vivo* (Tessler et al. 2020). In addition, we recombinantly expressed the putative luciferase gene from *P. atlanticum*, *pyroLuc*, and detected only low luciferase activity (Supplemental Figure S11). Specifically, the amount of light produced by PyroLuc is six orders of magnitude lower than RLuc, five orders of magnitude lower than AfLuc, and one order of magnitude lower than DafA. Similar to DafA, we were unable to detect enough luminescence to measure spectral emission for PyroLuc. Lastly, a later publication identified bioluminescent bacteria in the light-producing organs of *P. atlanticum* (Berger et al. 2021). For these reasons, the biochemical mechanism of bioluminescence in pyrosomes remains controversial and will benefit from future biochemical studies.

While most convergently evolved bioluminescence systems use non-homologous luciferases, there are several instances of parallel evolution (Delroisse et al. 2021) which provide intriguing insights into the factors that may shape the repeatability of molecular evolution. For example, fireflies and click beetles, members of the same order (Coleoptera) in the phylum Arthropoda evolved luciferases in parallel at least three times by recruiting members of the fatty acyl-coA synthetase gene family (He et al. 2024). These luciferases use ATP as a cofactor to adenylate D-luciferin, a luciferin substrate unique to fireflies and click beetles, which is then oxidized to produce light (McElroy et al. 1969). This pattern of evolution suggests that for D-luciferin based bioluminescence systems, luciferase evolution may be more repeatable, perhaps due to constraints imposed by the functional requirement of activating D-luciferin via adenylation and the widespread availability of the acyl-coA synthetase gene family for genetic recruitment (Karan et al. 2001). Unlike D-luciferin, coelenterazine does not need to be activated via biochemical modification, is used as a luciferin substrate across at least nine phyla, and reacts with a diversity of non-homologous luciferases to produce light, indicating the functional evolution of coelenterazine-based bioluminescence is not genetically constrained and often unrepeated. Deviating from this typical pattern of distinct genetic evolution, octocoral cnidarians and echinoderms each evolved luciferases by recruiting homologous haloalkane dehalogenases, a gene family with a sparse distribution across the tree of life as a result of horizontal transfer and subsequent gene loss. These findings underscore the role of historical contingency in shaping patterns of genetic recruitment during functional evolution. Divergent genetic histories, contingent on past mutational events, in combination with the number of potential genetic solutions, may explain when and why similar phenotypes evolve by recruiting similar versus distinct genes.

## Data availability

Protein sequences are available in GenBank: PP777633 (AfLuc), PP777634 (DafA), PP777635 (AF10707.1), PP777636 (AF17859.1), PP777637 (AF37282.1), PP777638 (AF37332.1), PP777639 (Gen224433), PP777640 (Uni20302.6), PP777641 (PyroLuc). Data files and code used to run analyses will be publicly available in Dryad following publication via the following link: https://doi.org/10.5061/dryad.rv15dv4gm.

## Acknowledgements

We thank Alexander Mikhailovsky for assisting with emission data collection and Vannie L Liu for assisting with recombinant protein expression. The work was supported by US National Science Foundation DEB-2153773 awarded to THO. The authors acknowledge use of Biological Nanostructures Laboratory (led by J. Smith) within the California NanoSystems Institute, supported by the UCSB and UCOP. This work was also supported by the Center for Scientific Computing (CSC), with computational facilities funded by the National Science Foundation (CNS-1725797). The CSC is supported by the California NanoSystems Institute and the Materials Research Science and Engineering Center (MRSEC; NSF DMR 1720256) at UC Santa Barbara. The work on this paper was supported by the Czech Science Foundation (GA22-09853S) and the Czech Ministry of Education, Youth and Sports (RECETOX RI LM2023069, e-INFRA LM2018140. This project was supported by the European Union’s Horizon 2020 research and innovation program under grant agreement No 857560 (CETOCOEN Excellence). This publication reflects only the author’s view, and the European Commission is not responsible for any use that may be made of the information it contains. DP is a Brno Ph.D. Talent Scholarship holder funded by the Brno City Municipality. J Delroisse is supported by an F.R.S.-FNRS research project (PDR, T.0071.23), previously held an F.R.S.-FNRS ‘Chargé de recherche’ fellowship (CR, 34761044), and also received financial support from an F.R.S.-FNRS research project (PDR, T.0169.20) and the Biosciences Research Institute of the University of Mons. WSB is PhD student under a FRIA fellowship (ID 40022483). ESL was funded by the National Science Foundation (GRFP 1650114). EP was supported by a Newton International Fellowship from the Royal Society (NIF\R1\222125). FM is supported by a Royal Society University Research Fellowship (URF\R1\191161) and a BBSRC research grant (BB/V01109X/1). NMH was funded by the National Science Foundation (PRFB 2011040). AM was funded by the National Institute of Health (R35-GM133530).

