## Supplemental Materials for "Functional characterization of luciferase in a brittle star indicates parallel evolution influenced by genomic availability of haloalkane dehalogenase"

### Supplemental Figures

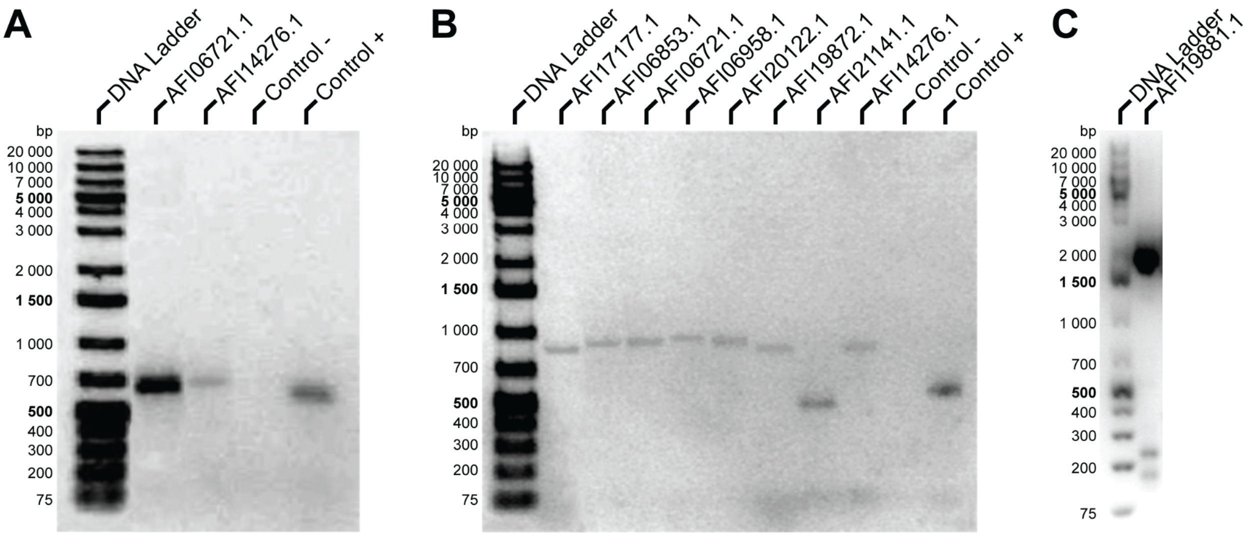


**Supplemental Figure S1. Results from genomic DNA-based validation PCRs.**

We performed PCRs to confirm portions of the HLD/LUC sequences. For all PCRs, we performed a negative control (absence of DNA) and positive control (amplification of a 600 bp fragment of the eukaryotic 18S ribosomal gene). All primer sets are detailed in Supplementary Table S2. (A) PCR amplification of AFI06721.1 and AFI14276.1 using primer set 1 produced single bands at the expected molecular weights. (B) PCR amplification of AFI17177.1, AFI06853.1, AFI06721.1, AFI06958.1 (*dafA*), AFI20122.1 (*afLuc*), AFI19872.1, AFI21141.1, and AFI14276.1 using primer set 2 produced single bands at the expected molecular weights. (C) PCR amplification of AFI19881.1 produced multiple bands — one at the expected molecular weight, and two other bands of lower molecular weight. However, these smaller bands did not impact downstream Sanger sequencing quality.

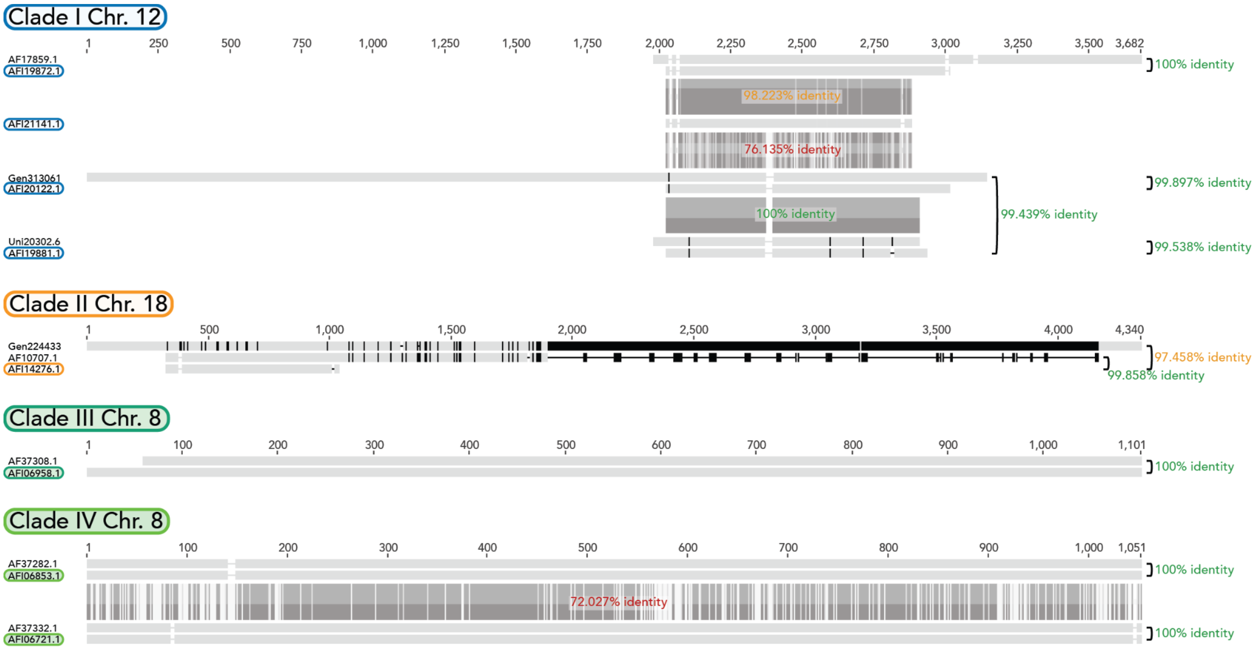


**Supplemental Figure S2. Multiple sequence alignment of the HLD/LUC gene models from Parey et al. (2024) and previous sequences from an earlier genome assembly and transcriptome.**

Gene models from the genome published in Parey et al. (2024) are highlighted and colored according to the chromosome in which they are found. Nucleotide differences between two synonymous sequences are denoted by black lines. Similarities between different pairs of gene models are denoted by gray lines.

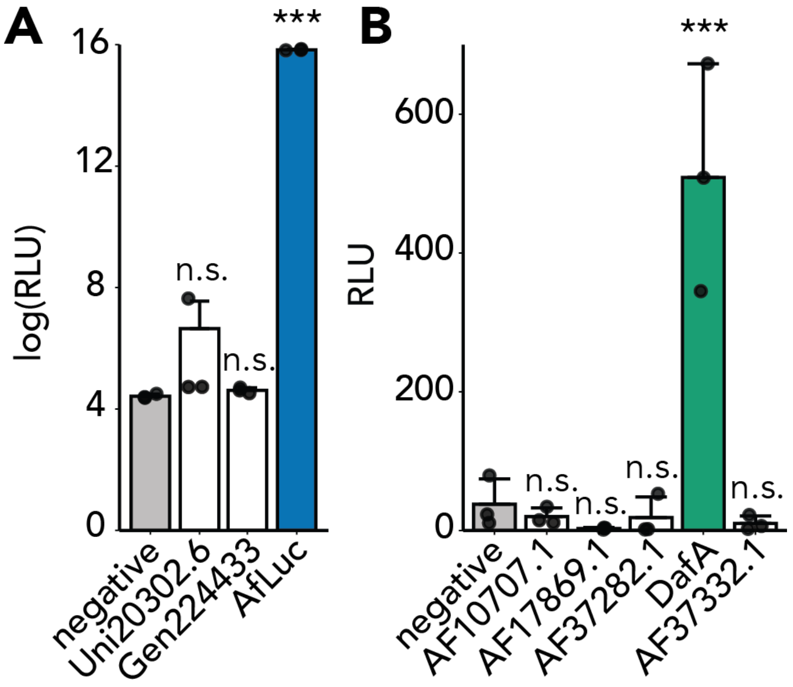


**Supplemental Figure S3. Crude extracts of AfLuc and DafA exhibit significant luciferase activity with coelenterazine.**

We recombinantly expressed dehalogenase genes from the genome of *Amphiura filiformis* and tested crude extracts for luciferase activity. (A) The negative control is a crude extract of bacteria transformed with an empty plasmid. (B) The negative control is a crude extract of bacteria expressing DhaA, a bacterial dehalogenase with no known luciferase activity. In both experiments, we tested for significant luciferase activity by using Dunnett's test (N = 3). Data in (A) are log-scaled. Data are expressed as average ± standard deviation.

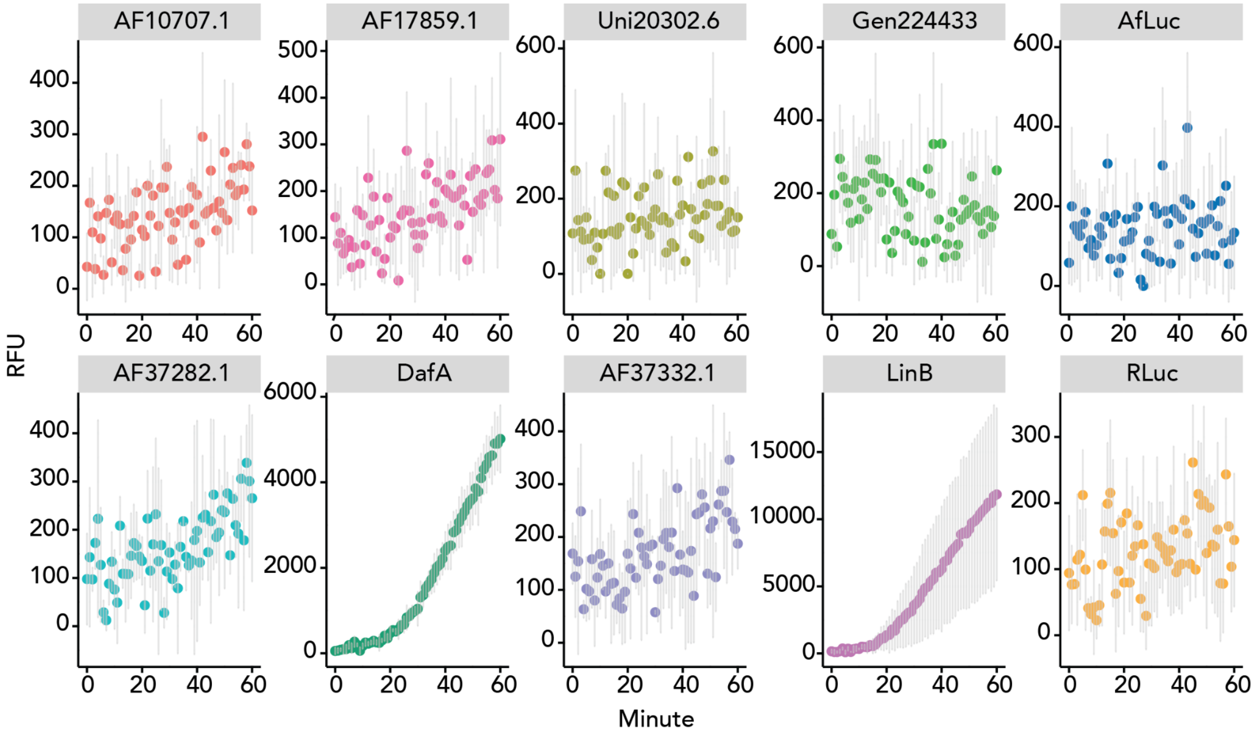


**Supplemental Figure S4. DafA exhibits dehalogenase activity with the substrate 1,2-dibromoethane.**

We recombinantly expressed each protein and tested crude cellular extracts for dehalogenase activity using RLuc as a negative control and LinB, a bacterial dehalogenase, as a positive control. We monitored reactions for 60 minutes. Data are expressed as average ± standard deviation (N = 3).

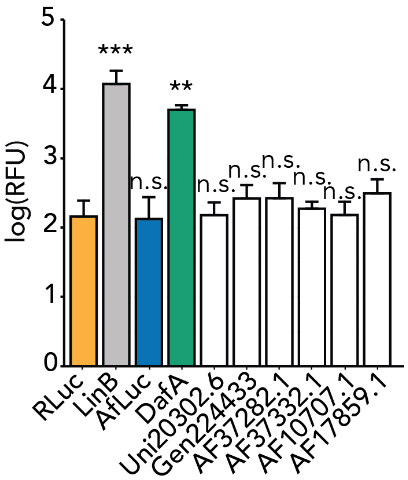


**Supplemental Figure S5. Only DafA exhibits significant dehalogenase activity with the substrate 1,2-dibromoethane.**

We compared the dehalogenase activity at 60 minutes for all recombinantly expressed dehalogenase sequences. Compared to the negative control of RLuc, only DafA significantly catalyzed the dehalogenation of 1,2-dibromoethane (Dunnett's test, $p = 0.00395$, $N = 3$). Data are log-scaled and expressed as average ± standard deviation.

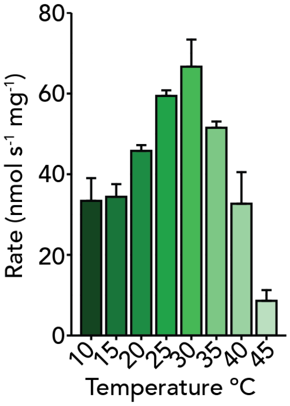


**Supplemental Figure S6. DafA's dehalogenase activity varies across temperatures.**

We measured DafA's dehalogenase activity with the substrate 1,2-dibromoethane across a range of temperatures. DafA exhibits maximum dehalogenase activity towards 1,2-dibromoethane at 30 ºC. The rate unit is nmol $s^{-1}$ $mg^{-1}$. Data are expressed as means ± standard deviation ($N = 3$).

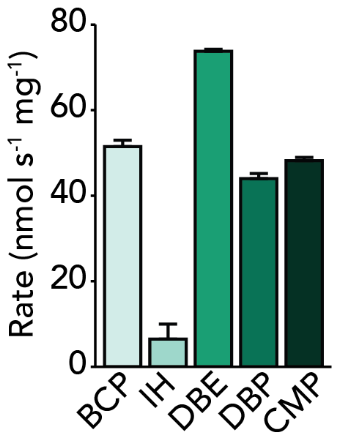


**Supplemental Figure S7. DafA catalyzes the dehalogenation of multiple halogenated substrates**

We tested DafA's ability to convert several halogenated substrates: (BCP = 1-bromo-3-chloropropane, IH = 1-iodohexane, DBE = 1,2-dibromoethane, DBP = 1,3-dibromopropane, CMP = 3-chloro-2-methylpropene). Rate is in nmol $s^{-1}$ $mg^{-1}$. For all plots, data are expressed as the average ± standard deviation.

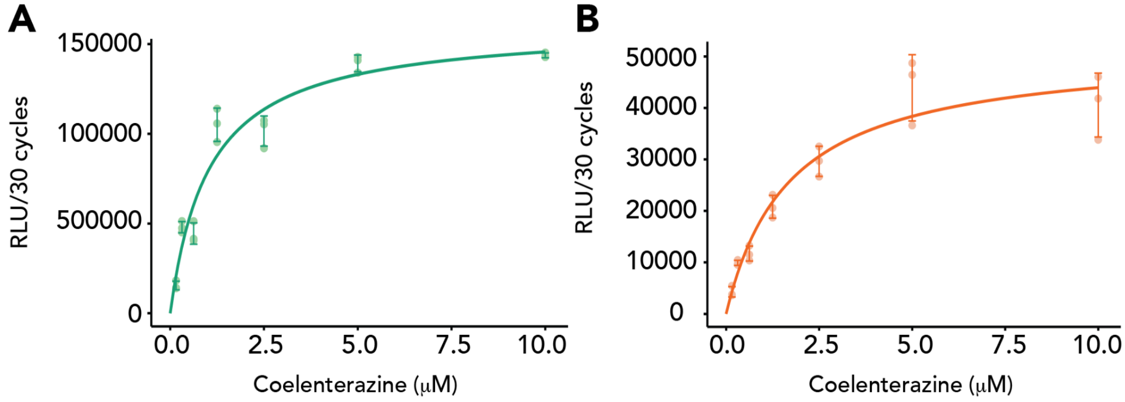


**Supplemental Figure S8. Michaelis-Menten kinetic profiles for DafA (green) and PyroLuc (orange).**

We varied the concentration of the coelenterazine substrate and measured light production for (A) DafA and (B) PyroLuc. For each measurement, we summed the light produced over 30 cycles (around 40 seconds).

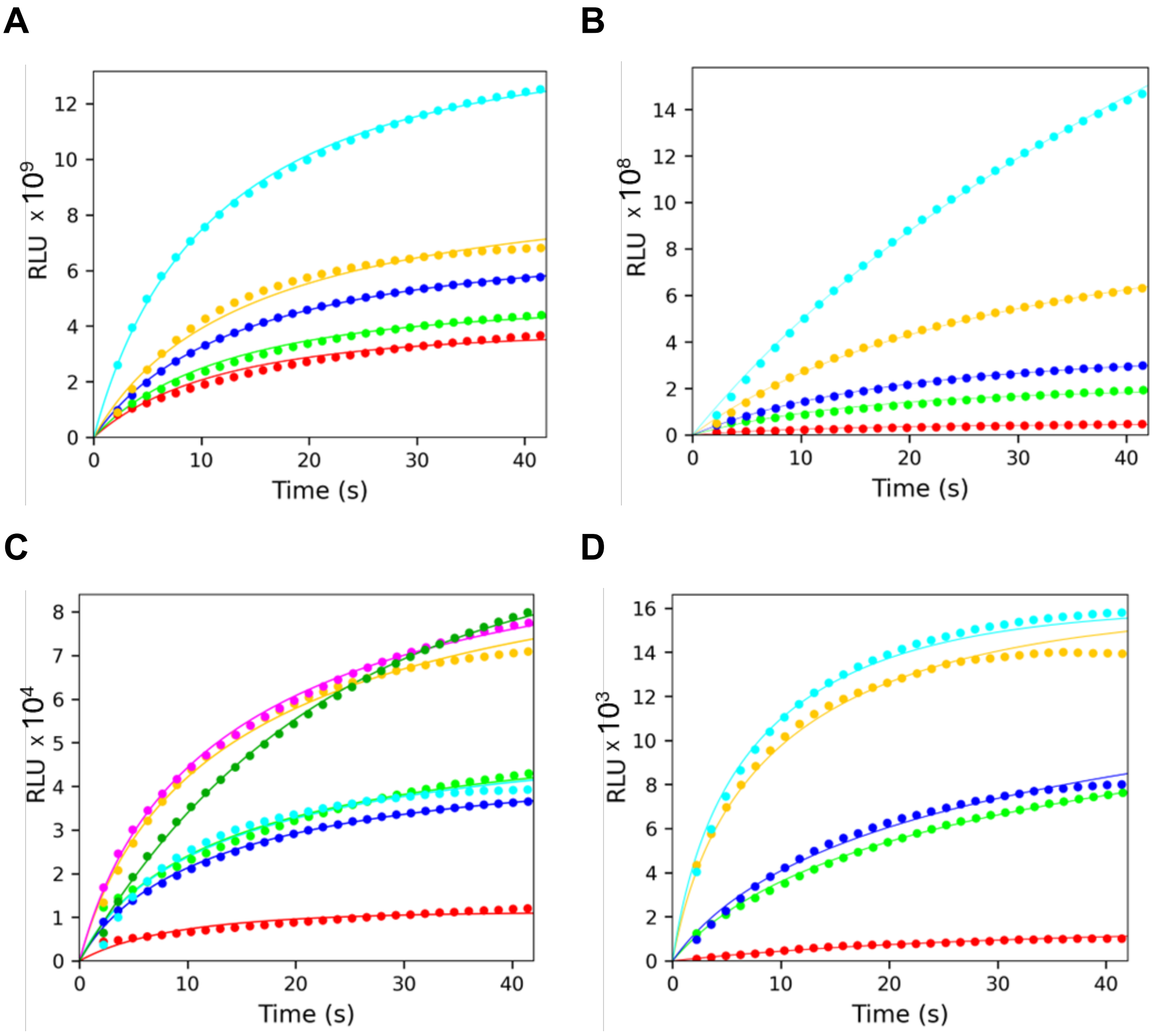


**Supplemental Figure S9. Steady-state kinetic data.**

Progress curves obtained by transformation of luminescence data recorded upon mixing 10 µM coelenterazine with 25 nM enzyme (final concentrations), (**A**) RLuc, (**B**) AfLuc, (**C**) DafA, and (**D**) PyroLuc. Each progress curve represents an average of 3 replicates. The solid lines represent the best fit.

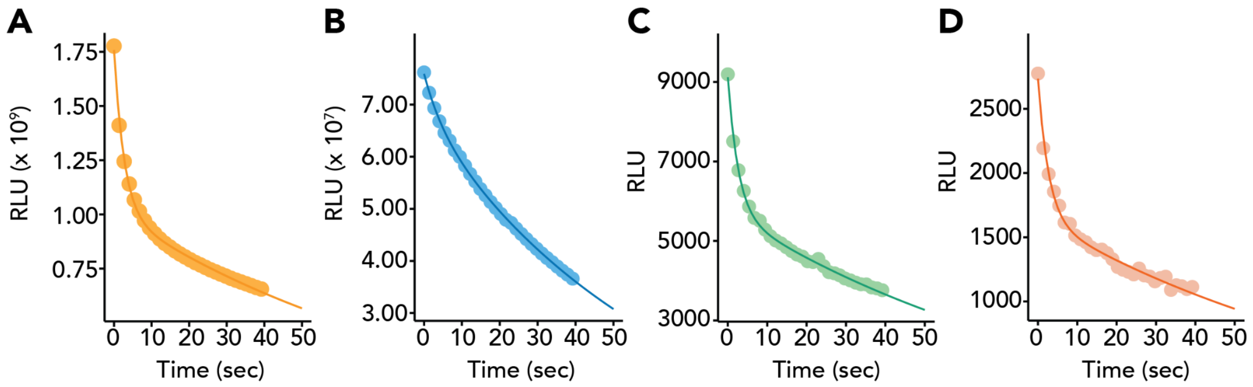


**Supplemental Figure S10. Luminescence decay data fit to a biexponential decay model.**

Results from estimating the luminescence decay parameters in (A) RLuc, (B) AfLuc, (C) DafA, and (D) PyroLuc. After combining 25 nM of recombinant protein with 10 μM coelenterazine, we recorded light production for around 40 seconds, averaged these data (N = 3), and fit them to a biexponential decay model.

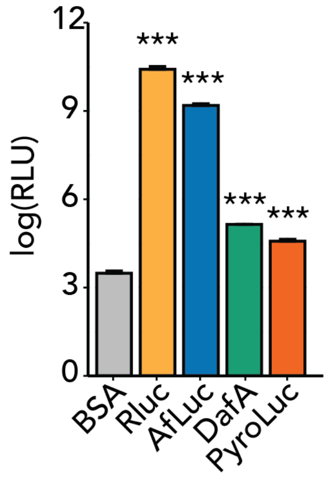


**Supplemental Figure S11. RLuc, AfLuc, DafA, and PyroLuc exhibit significant luciferase activity.**

Results of the luciferase assay show that RLuc (orange), AfLuc (blue), DafA (green), and PyroLuc (red) has significant luciferase activity with coelenterazine, relative to the negative control of BSA. We used Dunnett's test to assess significant luciferase activity (N = 3). Data are log scaled and expressed as means ± standard deviation.

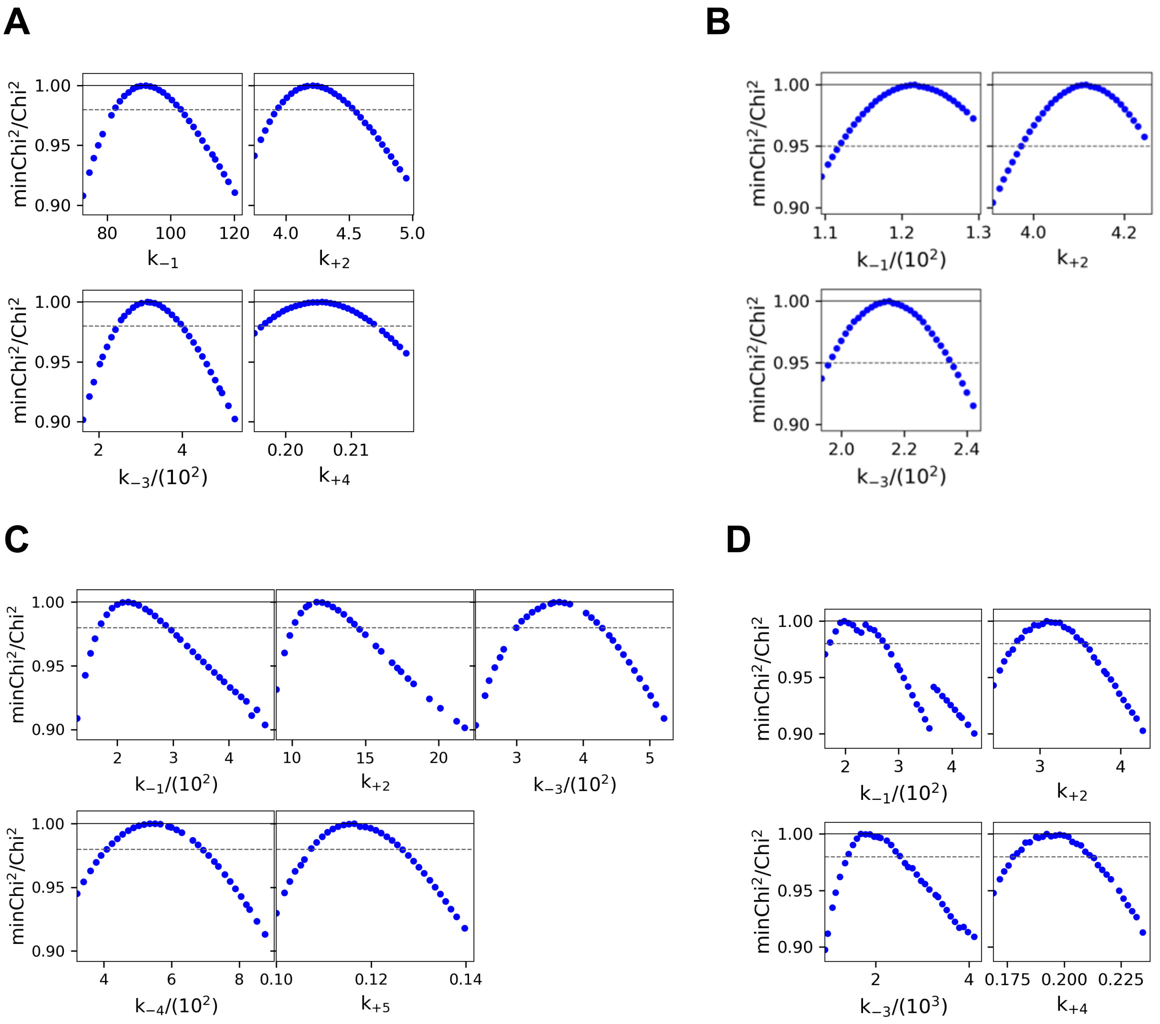


**Supplemental Figure S12. Confidence contour analysis.**

The confidence contours obtained for the steady-state parameters of (**A**) RLuc, (**B**) AfLuc, (**C**) DafA, and (**D**) PyroLuc. The gray dashed lines represent the $\chi^2$ threshold of 0.98, defining the lower and upper limits for each parameter presented in Supplementary Table S4.

### Supplemental Tables

| Name of sequences tested in this manuscript | Most similar gene model In Parey et al. (2024) |
|---|---|
| "*afLuc*", the sequence Gen313061 in Delroisse et al. (2017) | AFI20122.1 |
| "*dafA*", or preliminary gene model AF37308.1 | AFI06958.1 |
| AF17859.1 | AFI19872.1 |
| Uni20302.6 in Delroisse et al. (2017) | Ambiguous |
| AF10707.1 | AFI14276.1 |
| Gen224433 in Delroisse et al. (2017) | AFI14276.1 |
| AF37282.1 | AFI06853.1 |
| AF37332.1 | AFI06721.1 |

**Supplemental Table S1. Synonymized sequences of preliminary gene models from a draft genome of *Amphiura filiformis* with the final gene models of the genome published in Parey et al. (2024).**

We identified synonymous sequences based on percent sequence identity, as illustrated in Supplemental Figure S1.

| Gene | Primers (5’ → 3’) | Product (nt) | PCR |
|---|---|---|---|
| AFI17177.1 | *Forward* TTGGCACCAACTTCGAGCTC (set 2)<br>*Reverse* CCGATTTCAGCTGGGGAATCT (set 2) | 884 (set 2) | 1 band |
| AFI06853.1 | *Forward* ACCTTTCACTTGCACCGTCT (set 2)<br>*Reverse* ACAAATCCGACCGAAGCCAA (set 2) | 916 (set 2) | 1 band |
| AFI06721.1 | *Forward* GTCCATCCCACGTCAACCTC (set 1)<br>*Forward* AGTCCCTTCACTCTCACCGT (set 2)<br>*Reverse* TGCAAACGGTTCGCGATAAG (set 1)<br>*Reverse* GTCCCAACTTTCCGTAGCGT (set 2) | 627 (set 1)<br>911 (set 2) | 1 band |
| AFI06958.1 (*dafA*) | *Forward* AATCTCTGACAGCTTCCCCT (set 2)<br>*Reverse* GCACTCTGGCCGCATTAATG (set 2) | 956 (set 2) | 1 band |
| AFI20122.1 (*afLuc*) | *Forward* GCAAGCTTCTCCGGTCTCTT (set 2)<br>*Reverse* GGCAAAACCCCAGGACAAAC (set 2) | 925 (set 2) | 1 band |
| AFI19872.1 | *Forward* AGCTTCTCCACACTCTGTTGG (set 2)<br>*Reverse* CTCTCATCGGTGTAGGGTGC (set 2) | 859 (set 2) | 1 band |
| AFI21141.1 | *Forward* CTCTCATCGGTGTAGGGTGC (set 2)<br>*Reverse* TCAACACCAACGCTTCTCCA (set 2) | 494 (set 2) | 1 band |
| AFI19881.1 | *Forward* AGAACGGTCGCCAAAAGAGT<br>*Reverse* AAGCGGTGAGTTGCAGTGTA | 1604 (theoretically) | Multiple bands |
| AFI14276.1 | *Forward* CATTCCTCATGTTGCGCCTG (set 1)<br>*Forward* TGACACTCACCTTCTGCTGG (set 2)<br>*Reverse* TGACACTCACCTTCTGCTGG (set 1)<br>*Reverse* TACTGCGGACTCCCAACAAG (set 2) | 652 (set 1)<br>875 (set 2) | 1 band |

**Supplemental Table S2. Primer sequences used to amplify HLD/LUC sequences from the genome of *Amphiura filiformis*.**

| Substrate specificities ($nmol.s^{-1}.mg^{-1}$) | | | | |
|---|---|---|---|---|
| | DafA[1] | LinB[2] | DmmarA[3] | DnbA[4] |
| 1-iodohexane | 6.27 ± 3.50 | 22.02 ± 0.66 | 2.95 ± 0.07 | 1.05 ± 0.15 |
| 1,2-dibromoethane | 65.50 ± 6.91 | 84.74 ± 2.82 | 1.94 ± 0.13 | 7.83 ± 0.30 |
| 1,3-dibromopropane | 44.00 ± 1.19 | 63.26 ± 1.20 | 5.86 ± 0.00 | 6.48 ± 0.20 |
| 1-bromo-3-chloropropane | 37.07 ± 2.20 | 52.95 ± 1.58 | 5.16 ± 0.11 | 4.77 ± 0.06 |
| 3-chloro-2-methylpropene | 48.19 ± 0.78 | 43.34 ± 0.68 | 0.55 ± 0.17 | 1.35 ± 0.40 |

**Supplemental Table S3. Specific dehalogenase activities of DafA enzyme and other well-characterized haloalkane dehalogenases towards several halogenated substrates.**

DafA exhibits dehalogenase activity comparable to other characterized bacterial dehalogenases. The data are expressed as means ± the standard deviations.

[1] Data from this study

[2] Data were measured at 37 ºC and obtained in Buryska et al. (2019)

[3] Data were measured at 25 ºC and obtained in Vasina et al. (2022)

[4] Data were measured at 35 ºC and obtained in Vasina et al. (2022)

| | Rluc | Afluc | DafA | PyroLuc |
|---|---|---|---|---|
| $K_M$ (μM) | 0.91 ± 0.05 | 1.21 ± 0.03 | 2.0 ± 0.2 | 2.4 ± 0.3 |
| lower - upper limit | 0.83 - 1.03 | 1.12 -1.29 | 1.7 - 2.9 | 1.7 - 2.7 |
| $k_{cat}$ ($s^{-1}$) | 4.2 ± 0.2 | 4.12 ± 0.05 | 11 ± 1 | 3.0 ± 0.3 |
| lower - upper limit | 3.94 - 4.52 | 3.97 - 4.24 | 10.2 - 14.2 | 2.72 - 3.51 |
| $K_P$ (μM) | 0.32 ± 0.04 | 0.47 ± 0.02 | 0.29 ± 0.06 | 0.05 ± 0.01 |
| lower - upper limit | 0.32 - 2.22 | 0.43 - 0.51 | 0.15 - 0.24 | 0.04 - 0.07 |
| $K_{SI}$ (μM) | n.d. | n.d. | 5.9 ± 0.7 | n.d. |
| lower - upper limit | n.d. | n.d. | 4.2 - 6.8 | n.d. |
| $k_i$ ($s^{-1}$) | 0.21 ± 0.01 | n.d. | 0.12 ± 0.01 | 0.19 ± 0.01 |
| lower - upper limit | 0.20 - 0.21 | n.d. | 0.11 - 0.13 | 0.18 - 0.21 |

**Supplemental Table S4. Steady-state kinetic parameters.**

The estimated values ± standard errors calculated from the covariance matrix during nonlinear regression. The lower and upper limits for each parameter derived from the confidence contour at $\chi^2$ threshold 0.98 (Supplemental Figure S12); n.d. = not determined.

| | Lambda | Std error | p-value | Lambda_b | Std error | p-value | AICc |
|---|---|---|---|---|---|---|---|
| RLuc | 1.160e-02 | 3.453e-03 | 0.00117 | 3.728e-01 | 1.301e-01 | 0.00524 | 3685.06 |
| AfLuc | 2.061e-01 | 2.940e-01 | 0.485 | 1.576e-02 | 3.411e-03 | 1.33e-05 | 3084.84 |
| DafA | 1.125e-02 | 3.317e-04 | < 2e-16 | 3.800e-01 | 1.614e-02 | < 2e-16 | 1092.20 |
| PyroLuc | 1.120e-02 | 2.269e-03 | 3.86e-06 | 3.703e-01 | 9.584e-02 | 0.00216 | 1211.83 |

**Supplemental Table S5. Luminescence decay parameters estimated for RLuc, AfLuc, DafA, and PyroLuc.**

We performed non-linear model fitting and tested four different non-linear models. Based on AICc and residuals, the bi-exponential decay model was the best fitting model for all the datasets.

| Taxon | | Number of Origins of Coelenterazine Usage | Reference |
|---|---|---|---|
| Bacteria | | 0 | |
| Fungi | | 0 | |
| Radiolaria | Polycystine & Phaeodarian radiolarians | 2 | (Haddock et al. 2010) |
| Ctenophora | | 1 | (Haddock et al. 2010) |
| Porifera | | 1 | (Martini et al. 2020) |
| Octocorallia | | 1 | (DeLeo et al. 2024) |
| Hexacorallia | *Parazoanthus* | 1 | (Cormier et al. 1973) |
| | Hormathiidae | 1 | (Bessho-Uehara et al. 2020) |
| Medusozoa | Hydrozoa | 1 | (Shimomura 2019) |
| | *Periphylla periphylla* (Scyphozoa) | 1 | (Shimomura et al. 2001) |
| | *Pelagica noctiluca* (Scyphozoa) | 1 | (Morin and Hastings 1971) |
| Chaetognatha | *Eukrohnia fowleri* | 1 | (Thuesen et al. 2010) |
| | *Caecosagitta macrocephala* | 1 | (Thuesen et al. 2010) |
| Mollusca | *Vampyroteuthis infernalis* | 1 | (Robison et al. 2003) |
| | *Watasenia scintillans* | 1 | (Tsuji 2002) |
| | Ommastrephidae | 1 | (Takahashi and Isobe 1994; Galeazzo et al. 2019) |
| | *Pholas dactylus* | 1 | (Tanaka et al. 2009) |
| Arthropoda | *Conchoecia pseudodiscophora* (Halocyprid | 1 | (Oba et al. 2004) |

| | | | |
|---|---|---|---|
| | ostracods) | | |
| | Copepod | 1 | (Oba et al. 2009) |
| | *Oplophorus gracilirostris* (Decapoda) | 1 | (Shimomura et al. 1978) |
| | *Gnathophausia ingens* (Lophogastrida) | 1 | (Frank et al. 1984) |
| Chordata | *Oikopleura labradoriensis* | 1 | (Galt and Flood 1998) |
| | *Etmopterus* (Shark) | 1 | (Mizuno et al. 2021) |
| | Myctophidae (Myctophiformes) | 1 | (Duchatelet et al. 2019) |
| | *Vinciguerria attenuata* (Stomiiformes) | 1 | (Rees et al. 1990) |

**Supplemental Table S6. Number of origins of coelenterazine bioluminescence**

### Supplemental Materials and Methods

#### Materials

We purchased native coelenterazine (purity > 95 %, Catalog # 303-500) and NanoFuel Solvent (Catalog # 399) from Prolume LTD (Pinetop, AZ) and prepared single use aliquots by dissolving 500 µg of coelenterazine in 500 µL of nanofuel solvent to a stock concentration of 2.3615 mM, then stored the aliquots at -80 ºC. The pET21b(+)-Is-PETase was a gift from Gregg Beckham & Christopher Johnson (Addgene plasmid # 112202; http://n2t.net/addgene:112202 RRID:Addgene_112202). We purchased oligonucleotide primers and gBlocks™ from Integrated DNA Technologies (Coralville, IA). We purchased Q5 DNA polymerase (Catalog # M0491S), agarose gel extraction kit (Catalog # T1020S), reagents for Gibson assembly (Catalog # E2611L), competent *E. coli* 10-beta cells (Catalog # C3019H) for plasmid propagation, and competent *E. coli* BL21 cells (Catalog # C2527H) for protein expression from New England Biolabs (Ipswich, MA). We purchased reagents for extracting and purifying plasmids (Catalog # A1223) from Promega (Madison, WI) and reagents for Quick Start™ Bradford protein assay (Catalog # 5000205), the Bovine Serum Albumin (BSA) standard for calibration (Catalog # 500-0201), and Sodium dodecyl-sulfate polyacrylamide gel electrophoresis (SDS-PAGE) gels (Mini-PROTEAN TGX stain-free protein gels, Catalog # 4561094) from BioRad (Hercules, CA). We purchased Ni-NTA agarose (Catalog # R90115) and gravity-flow chromatography columns (Catalog # 34924) from Qiagen (Hilden, Germany). We purchased Amicon® Ultra centrifugal filter units with a 10 kDa molecular cutoff weight (Catalog # UFC901008) from Sigma Aldrich (St. Louis, MO).

#### Primer design, A. filiformis sampling, genomic DNA extraction and amplification of dehalogenase sequences

Based on the predicted HLD/LUC sequences of *A. filiformis*, several primer pairs were designed using the Primer3 software (v4.1.0, http://bioinfo.ut.ee/primer3). Primers were first *in silico* tested on the *A. filiformis* genome using the AmplifX software (v2.1.1 by Nicolas Jullien; Aix-Marseille Univ, CNRS, INP, Inst Neurophysiopathol, Marseille, France - https://inp.univ-amu.fr/en/amplifx-manage-test-and-design-your-primers-for-pcr). Primer pairs specifically amplifying a single target sequence were then selected. To confirm the sequences, validation PCRs were performed on genomic DNA of the brittle star. Due to its bacterial potential origin, these genes do not contain introns (Delroisse et al. 2017). Primer sequences are listed in Supplemental Table S2.

We collected *A. filiformis* (Müller, 1776) individuals with an Eckman grab at a depth of 30–40 m in the Gullmarsfjord near the Kristineberg Marine Research Station (University of Gothenburg, Fiskebäckskil, Sweden) in Summer 2023, rinsed sediments with fresh seawater to recover brittle stars, and maintained them in aquaria. We extracted genomic DNA from fresh brittle star arm fragments with the “DNeasy® Blood & Tissue” kit (QIAGEN), following the manufacturer protocol. We checked DNA integrity using gel electrophoresis, estimated concentration using a microspectrophotometer (DS-11 DeNovix Spectrophotometer), and stored DNA at -20 °C. We amplified sequences of interest using the “Red'y'Star Mix” commercial kit (Eurogentecs), following the manufacturer protocol. For each PCR, we performed a negative control (DNA replaced by PCR water) and a positive control (primers targeting the eukaryotic 18S ribosomal RNA gene). The PCR cycle was as follows: 10 min at 95 °C, followed by 35 cycles, with each cycle consisting of 20 s at 94 °C, 30 s at the primer hybridization temperature (as determined using the AmplifX program), and 1 min at 72 °C; with a final extension step of 10 min at 72 °C. If no amplification was possible using the “Red'Y'Star Mix”, we performed PCR using using the Q5® High-Fidelity DNA Polymerase (New England BioLabs) and a touchdown PCR protocol to increase PCR sensitivity and yield. The PCR cycle was as follows: 4 min at 95 °C, followed by 22 cycles, with each consisting of 30 s at 95 °C, 30 s at the primer hybridization temperature + 11 °C – decreased temperature by 0.5 °C every 1 cycle, 1 min – 1 min 30 at 72 °C, followed by 13 cycles consisting each of 30 s at 95 °C, 30 s at the primer hybridization temperature, 1 min – 1 min 30 at 72 °C, with a final extension step of 7 min at 72 °C).

We loaded PCR products onto an 1% agarose Tris-Borate-EDTA gel and performed electrophoresis to assess the molecular weights of each obtained band. After electrophoresis, we stained the gel in a TBE solution containing a GelRed intercalating agent (1:3000 dilution; GelRed® Nucleic Acid Gel Stain, Biotium) for 30 minutes under agitation, and visualized the gel using the Bio Rad Gel Doc2000 Imaging system to identify fluorescent bands. For PCR products containing a single band, we purified them by using the “SmartPure PCR Kit” DNA Purification Kit (Eurogentecs). For PCR products containing multiple bands, we excised the band corresponding to the expected molecular weight, melted the agarose by incubating for 10 min at 80°C, and used the resulting mixture as a DNA template for a second PCR. We measured the concentration of purified DNA using microspectrophotometry and performed Sanger sequencing (Eurofins Genomics, Germany). Then, we aligned these sequences with the reference HLD/LUC genes of *A. filiformis* to verify their identity (using MAFFT G-INS-I, implemented in Geneious Prime v2024.0.5, 2005, Dotmatics, USA).

**Cloning and expression of recombinant proteins**

We codon-optimized and synthesized DNA sequences corresponding to the luciferase sequence of *Renilla reniformis* (UniProt Accession P27652) and dehalogenase sequences from *Amphiura filiformis*, namely Gen224433, Gen313061 (named *afLuc*), and Uni20302.6 as reported in (Delroisse et al. 2017), and AF10707.1, AF17859.1, AF37282.1, AF37308.1 (named *dafA*), AF37332.1 (protein sequences predicted from a draft genome of *A. filiformis*). We cloned these sequences into the bacterial expression vector pET21b, a plasmid that encodes a T7 promoter, ampicillin resistance, and a C-terminus hexahistidine tag for purification. We transformed competent *E. coli* NEB® 10-beta cells via electroporation to propagate plasmids, extracted and miniprepped the plasmids, and sequenced the plasmids (Genewiz, South Plainfield, NJ) to confirm successful cloning. Then, we transformed competent BL21 cells via electroporation with these plasmids for protein expression. We grew up transformed BL21 cells in Terrific Broth containing ampicillin (at a final concentration of 100 µg/mL) at 37 ºC (shaking at 250 rpm) to mid-log phage (optical density at 600 nm ~ 0.4), then added isopropyl-β-D-thiogalactopyranoside (IPTG) to a final concentration of 1 mM to induce protein expression. We moved the bacterial cultures to a shaker at room temperature (250 rpm) and continued protein expression for 16-18 hours. We harvested bacterial cells by centrifuging cultures (4000 x g, 10 minutes) at 4 ºC, removed the supernatant, and froze the cell pellets. To lyse the bacterial cells, we resuspended the cell pellets in lysis buffer (10 mM TrisHCl, 500 mM NaCl, 0.5 % Tween20, 5 mM β-mercaptoethanol, pH 7.4) and sonicated the cells on ice (5 cycles of 30 s on, 10 s off at 30 % amplitude, MiSonix Sonicator 4000 with microtip probe). To pellet the cellular debris, we centrifuged the lysed cells (12,000 x g, 30 minutes) at 4 ºC, then collected the supernatant.

**Testing crude extracts of recombinant proteins for luciferase activity**

*Testing Gen224433, Gen313061, and Uni20302.6*

To test Gen224433, Gen313061, and Uni20302.6 for luciferase activity, we added 5 uL of 4 µM coelenterazine to 90 uL of the supernatant from lysed bacterial cells and measured luminescence using a microplate reader, set to an integration time of 3 seconds (Tecan Spark). As a negative control, we also tested an extract from BL21 cells transformed with an empty pQE80L plasmid. To test whether the expressed proteins exhibited luciferase assay, as compared to the negative control, we performed Dunnett's test (N = 3).

*Testing AF10707.1, AF17859.1, AF37282.1, AF37308.1, and AF37332.1*

To test AF10707.1, AF17859.1, AF37282.1, AF37308.1, and AF37332.1, we transferred single BL21(DE3) colonies of transformed cells into sterile 96-well plates (MTP) containing 100 µL of LB medium supplemented with ampicillin (100 µg/mL). We covered the plates with an air-pore membrane and cultivated for 3 h at 37 °C and 200 rpm. To induce the protein expression, we added 100 µl of LB medium with ampicillin (100 µg/mL) and IPTG (2 mM) to the mini-cultures and incubated MTP at 20 °C, 200 rpm, for 18 hours. We harvested cultures by centrifugation at 4 °C, 1600 g for 20 min, and froze the pellet immediately at -70 °C. After several hours we thawed the pellets at laboratory temperature for 10 min, and added 70 µL of a lysis buffer (20 mM potassium phosphate, 20 mM $Na_2SO_4$, and 1 mM EDTA, pH = 8.0) containing lysozyme (1 mg/mL) to each well. We incubated the MTP plate at 23 °C for 1 hour and removed cell debris from the lysate by centrifugation at 1600 g for 20 min. We transferred the supernatant into a new MTP and discarded the pellet. We used cell-free extracts from cells transformed with pet21b::DhaA and pet21b::RLuc as the negative and positive control, respectively. To test luciferase activity, we transferred 25 µL aliquots of cell-free extracts into a new MTP plate. We measured the baseline luminescence signal using FLUOstar Omega Microplate Reader (BMG labtech, Germany) for 5 s, with a gain set to 3200. Then, we added 225 µL of assay buffer (100 mM potassium phosphate, 1 mM $Na_2SO_4$, pH = 7.5) supplemented with 8.8 µM coelenterazine to the measured samples and immediately measured luminescence for 70 s. We performed all measurements in triplicates and then calculated the relative activity based on the peak area. We reported the results as averages along with their standard deviations. To test whether the expressed proteins exhibited luciferase assay, as compared to the negative control, we performed Dunnett's test (N = 3).

**Recombinant expression and purification of AfLuc, RLuc, DafA, and PyroLuc**

We grew up BL21 cells, transformed with plasmids encoding RLuc, AfLuc, DafA, and PyroLuc, in 50 mL of Terrific Broth (containing ampicillin at a final concentration of 100 µg/mL) at 37 ºC (shaking at 250 rpm). To induce protein expression, we added 50 µL of IPTG to a final concentration of 1 mM. We moved the bacterial cultures to room temperature and continued shaking at 250 rpm for 18 hours. To harvest the bacterial cells, we centrifuged the cultures (4,000 x g, 10 minutes) at 4 ºC, removed the supernatant, and froze the cell pellets at -20 ºC for 3 hours. We extracted recombinant protein by lysing bacterial cells in 5 mL of lysis buffer (10 mM TrisHCl, 500 mM NaCl, 30 mM imidazole, 0.5 % Tween20, 5 mM β-mercaptoethanol, pH 7.4), then sonicated the cells on ice (5 cycles of 30 s on, 10 s off at 30 % amplitude, MiSonix Sonicator 4000 with microtip probe), and centrifuged the crude lysate (12,000 x g, 30 minutes) at 4 ºC to pellet cellular debris. To purify the proteins, we collected the clarified supernatants, added it to 1 mL of Ni-NTA agarose beads, then mixed and incubated the samples at 4 ºC overnight on a rotary mixer. Next, we loaded the Ni-NTA slurry into gravity-flow chromatography columns, allowed the slurry to settle, then removed the cap and discarded the flow through. We washed the column twice, each time with 2 mL of wash buffer (10 mM TrisHCl, 500 mM NaCl, 30 mM imidazole, pH 7.4), and eluted proteins bound to the Ni-NTA agarose by adding 4 mL of elution buffer (10 mM TrisHCl, 500 mM NaCl, 500 mM imidazole, pH 7.4). We performed spin ultrafiltration, using centrifugal ultrafiltration units with a 10 kDa molecular weight cutoff, to concentrate and buffer exchange the eluates into a storage buffer (10 mM TrisHCl, 300 mM NaCl, 20 % glycerol). We ran SDS-PAGE to assess protein purity and performed a Bradford assay (Bio-Rad Laboratories, Inc.), calibrated by using a standard curve of BSA, to quantify yield of recombinant proteins. We flash froze single-use aliquots of recombinant protein and stored them in - 80 ºC until use.

**Collecting data for estimating Michaelis-Menten kinetics profiles**

We characterized the enzyme kinetics of AfLuc, DafA, and PyroLuc, using RLuc as a positive control and BSA and luciferin as negative controls. We added 50 μL of recombinant protein to 50 μL of coelenterazine, resulting in a total volume of 100 μL and final concentrations of 25 nM recombinant protein and varying concentrations of coelenterazine (10 μM, 5 μM, 2.5 μM, 1.25 μM, 0.625 μM, 0.3125 μM, 0.15625 μM). To perform the measurements, we used a plate reader (Tecan Spark) and first measured the background for 5 cycles, then injected coelenterazine and measured luminescence for 30 cycles, with a 1 second integration time for each cycle. We subtracted the background luminescence from each of the measurements after coelenterazine was injected. We repeated each sample measurement in triplicate.

**Kinetic data analysis and statistics**

An updated protocol applying new standards for collecting and fitting steady-state kinetic data (Johnson 2019; Schenkmayerova et al. 2021) was used to analyze and compare kinetic behavior of RLuc, AfLuc, PyroLuc and DafA. The recorded luminescence traces (rate vs. time) were transformed to reaction progress curves corresponding to cumulative luminescence in time (Supplemental Figure S8). The transformed kinetic data (product vs. time) were fitted globally with the KinTek Explorer (KinTek Corporation, USA). The software allows for the input of a given kinetic model via a simple text description, and the program then derives the differential equations needed for numerical integration automatically. Numerical integration of rate equations searching a set of kinetic parameters that produce a minimum $\chi^2$ value was performed using the Bulirsch–Stoer algorithm with adaptive step size, and nonlinear regression to fit data was based on the Levenberg–Marquardt method (Johnson et al. 2009a). To account for fluctuations in experimental data, enzyme or substrate concentrations were slightly adjusted (± 10%) to derive best fits. Residuals were normalized by sigma value for each data point. The standard error (S.E.) was calculated from the covariance matrix during nonlinear regression. In addition to S.E. values, more rigorous analysis of the variation of the kinetic parameters was accomplished by confidence contour analysis by using FitSpace Explorer (KinTek Corporation, USA). In these analyses (Supplemental Figure S12), the lower and upper limits for each parameter were derived (Supplemental Table S4) from the confidence contour obtained from setting $\chi^2$ threshold at 0.98 (Johnson et al. 2009b). The scaling factor, relating luminescence signal to product concentration, was applied as one of the fitted parameters, well constrained by end-point levels of kinetic traces recorded at particular substrate concentrations. The steady-state model (Supplementary Scheme S1) was used to obtain the values of turnover number $k_{cat}$, Michaelis constant $K_M$ and equilibrium dissociation constant for enzyme-product complex $K_p$. A conservative estimate for diffusion-limited substrate and product binding $k_1$ and $k_{-5}$ (100 µM$^{-1}$.s$^{-1}$) was used as a fixed value to mimic rapid equilibrium assumption. The basic kinetic model was extended by irreversible inactivation (significant for all enzymes except AfLuc) and substrate inhibition (indicated for DafA).

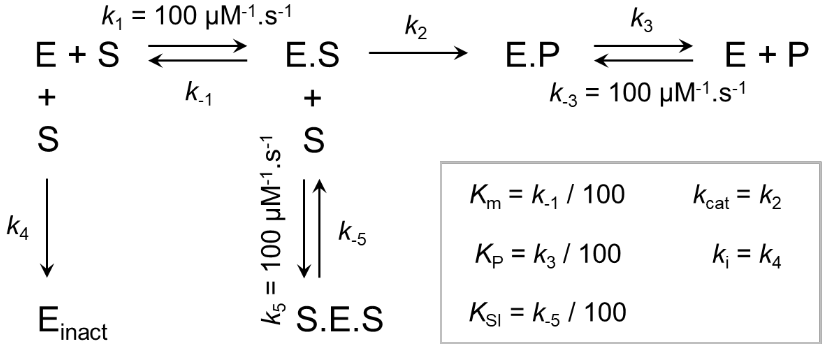


**Supplementary Scheme S1.** Steady-state model used for kinetic data analysis.

**Measuring luminescence emission spectra**

We measured emission spectra using a custom spectroradiometer set-up at UCSB, as detailed in a previous study (Hensley et al. 2021). In brief, we added 10 µL of 100 µM coelenterazine to 200 µL of purified recombinant proteins, diluted in 1X TBS pH 7.4 (final concentrations 4.76 µM coelenterazine with 1.17 µM Renilla luciferase, 0.68 µM AfLuc, 2.06 µM DafA, or 0.11 µM PyroLuc), and measured the emission spectra using a spectroradiometer (Acton SpectraPro 300i) with a charge-coupled device camera detector (Andor iDus). We corrected these spectral data using correction factors calculated from the spectrum of a black body-like light source (Ocean Optics LS-1) and subtracted background emission spectra data of 1X TBS buffer from the experimental data. We repeated each sample measurement in triplicate, then normalized and averaged these data.

**Testing whole cell extracts for dehalogenase activity**

To prepare whole cell extracts, we transferred transformed BL21(DE3) cells into sterile 96-well microtiter plates (MTPs) containing 100 µL of LB medium supplemented with ampicillin (100 µg/ml). We covered MTPs using a air-pore membrane and incubated them for 3 hours at 37 °C with shaking at 200 rpm. To induce protein expression, we added 100 µL of LB with ampicillin (100 µg/ml) and IPTG (2 mM) to each mini-culture and incubated MTPs at 20 °C for 18 hours, while shaking plates at 200 rpm. We harvested cell cultures by centrifuging MTPs (1,600 x g, 4 ºC, 20 minutes), and discarded the supernatant. To wash the cell pellets, we added 200 µL of reaction buffer (1 mM orthovanadate, 20 mM phosphate buffer, pH = 8.0), centrifuged MTPs (1,600 x g, 4 ºC, 20 minutes), and repeated this step again. To lyse cell pellets, we resuspended cell pellets in 200 µL of the reaction buffer and froze them at - 70 ºC.

To screen whole cell extracts for dehalogenase activity, we used a whole-cell halide oxidation (HOX) assay (Aslan-Üzel et al. 2020). We prepared the substrate 1,2-dibromoethane (DBE) in the reaction buffer (final concentrations 0.3 mM DBE, 1 mM orthovanadate, 20 mM phosphate buffer, pH = 8.0). In a new black bottom 96-well MTP, we added the assay master mix, which consists of 25 µM aminophenyl fluorescein, 26 mM $H_2O_2$, 1.1 U *Curvularia inaequalis* histidine-tagged vanadium chloroperoxidase, 1 mM orthovanadate, 20 mM phosphate buffer, pH = 8.0. We then added resuspended cells to each well for a final OD600 ~ 0.02, followed by 100 µL of DBE substrate to a final volume of 200 µL, and covered the plate. We immediately measured fluorescence with an excitation at 488 nm and emission detention at 525 nm, at 30 ºC (Synergy™ H4 Hybrid Microplate Reader), with results normalized to OD600 = 1. We measured all data in triplicate and calculated means and standard deviations.

**Measurements of specific dehalogenase activity at varying temperatures**

We measured the temperature profile and dehalogenase activity with various substrates for DafA using the capillary-based droplet microfluidic platform MicroPEX (Vasina et al. 2022), which enables us to measure specific enzyme activity within droplets for multiple enzyme variants in one run. We generated droplets (Mitos Dropix) and used a microfluidic pump to generate a custom sequence of droplets (150 nl aqueous phase, 300 nl oil spacing). These droplets are then incubated with the halogenated substrate via microdialysis and partitioning between the oil (FC 40) and the aqueous phase. This reaction solution consists of 1 mM HEPES, 20 mM $Na_2SO_4$, pH 8.2, and a complementary fluorescent indicator 8-hydroxypyrene-1,3,6-trisulfonic acid (50 µM HPTS). Then, fluorescence is measured using an optical setup with an excitation laser (450 nm), a dichroic mirror with a cut-off at 490 nm filtering the excitation light, and a Si-detector. Using this pH-based fluorescence assay, we were able to monitor enzymatic activity over four minutes. We processed raw data using LabView and used MatLab to calculate specific activities.
